# Preventing Microglial Reactivity Protects from Acute and Progressive Neuronal Dysfunction, Motor Impairments and Sedation following Alcohol Abuse

**DOI:** 10.1101/2025.06.12.658437

**Authors:** Evi Paouri, Sarah Stanko, Nadia Gasmi, Gabriele Glusauskas, Valery Mao, Andrew Kwak, Sahar Sarmadi, Emily Huang, Courtney Hershberger, Quinn Watercutter, Hope Burton, Alexis Crockett, Anxhela Gjojdeshi, Megan R. McMullen, Richa M. Hanamsagar, Daniel M. Rotroff, Richard A. Prayson, Staci D. Bilbo, Saba Valadkhan, Hod Dana, Laura E. Nagy, Dimitrios Davalos

**Affiliations:** Department of Neurosciences, Cleveland Clinic Research, Cleveland, OH 44106, USA; Department of Molecular Medicine, Cleveland Clinic Lerner College of Medicine of Case Western Reserve University, Cleveland, OH 44106, USA; Department of Inflammation and Immunity, Cleveland Clinic Research, Cleveland, OH; Department of Quantitative Health Sciences, Cleveland Clinic Research, Cleveland, OH, 44195, USA; Center for Quantitative Metabolic Research, Cleveland Clinic, Cleveland, OH, 44195, USA; Department of Pediatrics, Lurie Center for Autism, Massachusetts General Hospital for Children, Boston, MA 02421, USA; Endocrinology and Metabolism Institute, Cleveland Clinic, Cleveland, OH, 44195, USA; Division of Pathology, Cleveland Clinic, Cleveland, OH 44106, USA; Department of Psychology and Neuroscience, Duke University, Durham, NC, USA; Department of Neurobiology, Duke University Medical Center, Durham, NC, USA; Department of Molecular Biology and Microbiology, Case Western Reserve University, Cleveland, OH; Department of Biochemistry, Case Western Reserve University School of Medicine, Cleveland, OH 44106, USA

**Author notes:** These authors contributed equally to this work (Evi Paouri, Sarah Stanko).

## Abstract

Alcohol abuse is the primary risk factor for alcohol use disorder (AUD), a leading cause of preventable morbidity and mortality, characterized by systemic inflammation, multi-organ damage, and neurological impairments. While direct effects of alcohol on brain function are well-established, the role of microglia in acute and chronic neurological dysfunction in AUD remains unclear. Using longitudinal *in vivo* imaging in mice during acute and repeated alcohol abuse, we found that microglia exhibit dynamic morphological responses that precede but parallel ethanol-induced sedation. Ethanol also induced microglia-dependent synapse elimination and reduced neuronal activity and density. Genetic disruption of microglial MyD88 reversed these ethanol-associated changes in microglial reactivity, neuronal structure, and function, while protecting against alcohol-induced intoxication and motor impairments. These findings identify microglia as cellular drivers of acute and chronic brain dysfunction following alcohol abuse, and highlight MyD88 as a critical therapeutic target for the detrimental neurological consequences of AUD.

## Introduction

Alcohol use disorder (AUD) represents a critical public health crisis, affecting approximately 1 in 9 American adults (10.9%), yet the alcohol abuse behaviors that drive AUD development are far more widespread, with one in four adults in the United States reporting binge drinking in the past month^1^. The emergence of high-intensity drinking – consuming 8+ drinks for women, 10+ drinks for men within 2 hours, or twice the binge threshold – has created an even more dangerous pattern that peaks around age 21^1, 2^. While extensive research has focused on neuronal network changes underlying AUD—reward circuit dysfunctions associated with addiction, withdrawal symptoms, compulsive alcohol-seeking behaviors^3^—the neurotoxic mechanisms by which alcohol abuse impairs brain structure and function, in the much larger population engaging in binge and high-intensity drinking patterns, remain poorly understood. Yet, there were approximately 178,000 annual deaths from excessive alcohol use between 2020-2021, with one-third resulting from binge drinking patterns^1^, highlighting the urgent need to understand how alcohol abuse damages the brain in the millions of people who engage in these dangerous drinking behaviors.

Alcohol abuse including binge drinking and high-intensity drinking, induces severe systemic complications (such as cardiovascular, gastrointestinal, liver, pancreatic, musculoskeletal damage)^4, 5^, and immediate and progressive neurological impairments in humans and animal models^6–8^. Mechanistically, alcohol consumption impairs gut barrier integrity, releasing lipopolysaccharides (LPS) from gut bacteria into the bloodstream. Ethanol and LPS activate Toll-like receptors (TLRs), leading to liver inflammation and increased circulating inflammatory cytokines that promote neuroinflammation and neuronal damage^4, 6, 7, 9^. Additionally, ethanol readily crosses the blood-brain barrier and directly reaches brain cells, many of which express receptors that can be modulated by ethanol^7, 10–12^. Chronic alcohol use results in neuronal damage manifested as decreased dendritic arborization^13^, neuronal loss^14^ and reduced brain electrical activity and gray matter volume^15–18^. The frontal lobes, critical for decision making, impulse control, and emotional regulation, are particularly vulnerable^19–22^. Beyond its direct neurotoxicity, alcohol causes nutritional, metabolic, and vascular disruptions^8, 23, 24^, that, along with peripheral and brain-derived inflammation, orchestrate its detrimental impact on brain structure and function through mechanisms that have yet-to-be deciphered.

Microglia survey the brain by continuously scanning their microenvironment with highly motile processes that rapidly respond to tissue damage^25, 26^. Microglia contact dendritic spines constitutively or in an activity-dependent manner^27–32^, and can influence neural networks through synaptic pruning^28,33^ and release of neuroactive factors^34^. Microglia can also alter neuronal activity or respond to both neuronal hyperexcitability and silencing, as a result of general anesthesia^35, 36^ or experimental manipulations of neurons or microglia^31, 37, 38^. *In vivo* imaging showed that ethanol exposure during development had no impact on microglial tissue surveillance during adolescence^39, 40^, but ethanol acutely reduced it in the adult visual cortex^41^. Yet how acute or repeated alcohol abuse may affect microglial responses and whether those responses impact neuronal integrity and function, or animal behavior, remain unexplored.

Signaling through TLR4 and its adaptor protein myeloid differentiation factor 88 (MyD88) recruits protein kinases and transcription factors such as NF-κB, leading to upregulation of immune response genes^42^. TLR4 and MyD88 are essential for ethanol-induced inflammatory cytokine production in the mouse brain^43^, but also contribute to alcohol-induced sedation and motor impairments^44^. Microglia are major expressors of TLR4/MyD88 in the brain and show morphological changes indicative of inflammatory activation upon ethanol challenge both *in vitro* and in mouse and human brain tissue^10, 45–48^. RNA-sequencing (RNAseq) in prefrontal cortical microglia from alcohol exposure mouse models found upregulation of immune response genes, including TLRs^49, 50^. These findings point towards microglia and the TLR/MyD88 signaling pathway as neuroimmune contributors to AUD. However, it remains unclear whether microglia simply react to body-wide inflammation via this pathway, or actively use it to mediate alcohol’s effects on the brain.

Here, we investigated how alcohol abuse affects microglial responses and brain function in adult mice. To model binge and high-intensity drinking patterns in humans, we repeatedly administered either ataxia- or sedation-inducing ethanol doses, respectively. By tracking individual cells over 10 consecutive days, we found that microglia underwent gradual and partially reversible morphological changes, that preceded the onset of, yet paralleled the recovery from, each ethanol-induced sedation. Microglia also exhibited progressive worsening of their morphological reactivity following repeated sedative doses of ethanol, accompanied by a phagocytosis, matrisome and cell growth/adhesion transcriptional signature. These ethanol-induced microglial responses were accompanied by structural changes of the neuronal network including microglia-mediated elimination of synaptic structures, and dose-dependent reduction in neuronal cell density. Importantly, genetic deletion of *MyD88* in microglia fully reversed both acute and repeated/chronic effects of alcohol, preventing aberrant synapse loss, and neuronal cell death even after the entire 10-day course of alcohol abuse. Disruption of MyD88 signaling in microglia also protected mice from acute decreases in neuronal activity *in vivo* and allowed them to experience significantly reduced alcohol-induced sedation and motor impairments.

## Results

### Binge ethanol consumption induces acute and progressive morphological microglial responses *in vivo*

Microglia undergo pronounced morphological changes in response to a wide range of challenges that often reflect transitions in their molecular and functional states^51^. To begin unraveling how alcohol abuse may affect microglia, we implemented a model of binge drinking^52^, previously reported to cause microglial inflammatory activation, ROS production, and neurodegeneration in chronically treated mice^53^. Adult mice were orally administered 5g ethanol/kg of body weight (BW) once daily for 10 consecutive days and brains were harvested 5 hours after the last administration for histological evaluation using high-resolution confocal microscopy (Fig. 1a_i_). Three-dimensional (3D) morphological analyses showed reduced microglial process volume, length and number of branch points, and enlarged cell bodies in the somatosensory cortex of mice that underwent the 10-day ethanol course, compared to saline treated controls (Supp. Fig. 1a,b). Although the effects of this chronic binge drinking model on brain inflammation were established in male mice^54, 55^, we found no significant differences in microglial process complexity and soma volume between male and female animals (Supp. Fig. 1b).

**Fig. 1.**
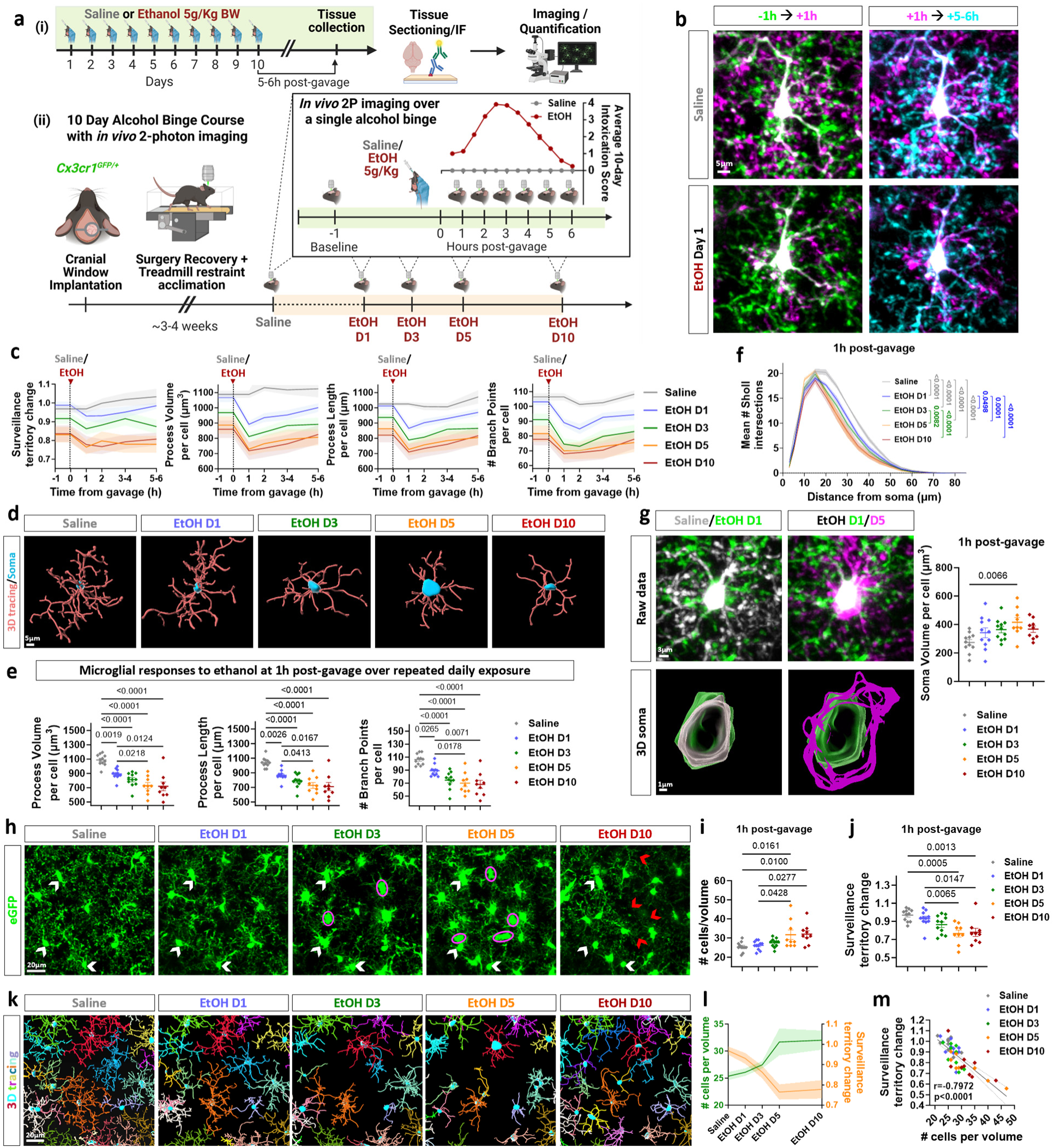
Binge ethanol consumption induces acute and progressive morphological microglial responses *in vivo.* **a.** Experimental design schematics. **(i)** Chronic binge drinking mouse model: daily ethanol treatment for 10 consecutive days (5 g/kg BW by oral gavage), followed by tissue collection and histological analyses. **(ii)** I*n vivo* two-photon microscopy experimental plan for imaging somatosensory cortical microglia in *Cx3cr1^GFP/+^*mice. This design allowed recording the evolution of microglial responses, following saline treatment and during onset and progression of acute ethanol-induced sedation (intoxication scoring curve shown in inset), as well as throughout the 10-day chronic binge ethanol exposure course, in the same animals. **b.** Representative two-photon microscopy images of the same microglial cell, captured at -1 h, +1 h and +5–6 h relative to gavage on saline treatment day and on ethanol day 1. Scale bar, 5 µm. **c.** Time course changes of microglial territory, process volume, process length, and number of branch points over hours following gavage on saline day and on binge ethanol exposure days 1, 3, 5, and 10. **d.** Representative 3D tracings of microglial processes and cell soma reconstructions from the same cell at 1 h post-gavage on saline day and binge ethanol exposure days 1, 3, 5, and 10. Scale bar, 5 µm. **e.** Quantification of microglial process volume, process length, and branch points at 1 h post-gavage on saline day and binge ethanol exposure days 1, 3, 5, and 10. **f.** Sholl analysis of microglia at 1h post-gavage on saline day and binge ethanol exposure days 1, 3, 5, and 10. Mixed-effects model with Geisser-Greenhouse correction and Tukey’s post hoc test. **g.** Representative two-photon microscopy images of the same microglial cell and its 3D-reconstructed cell soma, shown at 1 h post-gavage on saline treatment day and on binge ethanol exposure days 1 and 5. Scale bars, 3 µm (raw data), 1 µm (3D soma). Graph shows the quantification of cell soma volume at 1 h post-gavage on saline day and on binge ethanol exposure days 1, 3, 5, and 10. **h.** Representative two-photon microscopy images of microglia in the same cortical location at 1 h post-gavage on saline treatment day and on binge ethanol exposure days 1, 3, 5, and 10. White arrowheads indicate cells consistently tracked across all imaging days. Red arrowheads mark locations where cells were absent on ethanol day 10 compared to day 5. Magenta ellipses highlight new cells detected on ethanol days 3 and 5. Scale bar, 20 µm. **i-j.** Quantification of microglial cell number per analyzed volume (**i**), and surveillance territory change (**j**) at 1 h post-gavage on saline day and on binge ethanol exposure days 1, 3, 5, and 10. **k.** Representative 3D tracings of microglial processes and cell soma reconstructions from the same cortical volume imaged at 1 h post-gavage on saline day and on binge ethanol exposure days 1, 3, 5, and 10, showing progressive changes in surveillance territory. Scale bar, 20 µm. **l.** Time course changes in microglial cell number per analyzed volume overlaid with corresponding changes in surveillance territory throughout saline day and binge ethanol exposure days 1, 3, 5, and 10. **m.** Pearson correlation analysis showing the relationship between surveillance territory change and microglial cell number per imaged volume throughout the 10-day binge ethanol course. Data points are color-coded per treatment condition/timepoint. Data presented as mean ± SEM, and acquired from n=11 imaged cortical volumes, from 6 mice. **e, g, i-j.** One-way ANOVA with Tukey’s multiple comparison test. Only statistically significant comparisons are shown.

Having established that repeated episodes of binge-like drinking culminated in significant microglial phenotypical changes, we sought to better understand the evolution of their responses from the acute to the chronic stages of binge ethanol exposure. We thus performed *in vivo* imaging of microglia in awake microglial-reporter mice (*Cx3cr1^GFP/+^*) bearing cranial windows over somatosensory cortex, using two-photon microscopy. We imaged two brain volumes per animal, first after saline administration and then again on days 1, 3, 5, and 10 of the binge ethanol course (Fig. 1a_ii_). On each imaging day, baseline timelapse series were acquired 30-60 minutes before oral gavage, and each volume was re-imaged hourly for 5-6 hours post-gavage. This imaging period overlapped with the animals’ daily ethanol-induced sedation, which was scored using a modified alcohol intoxication scoring system^56^ (Fig. 1a_ii_, Supp. Fig. 1c). We then performed detailed morphological analyses of microglial processes and cell bodies in all longitudinally recorded 3D volumes (Supp. Fig. 2a). Notably, the significant loss of microglial ramification and surveillance territory in response to binge ethanol treatment peaked within approximately 1 hour following each ethanol dose –across both initial and subsequent days–preceding the period of peak sedation by 1-2 hours. These microglial morphological metrics showed minimal changes during saline treatment, when the animals remained alert (Fig. 1b-c, Supp. Fig. 2a-b). 3D tracing of microglial processes (Fig. 1d) showed a progressive reduction of process volume, length, branch points and Sholl intersections with repeated binge ethanol exposure, peaking on day 5 (Fig. 1e-f). These differences were less prominent 5-6 hours post-gavage (Fig. 1c, Supp. Fig. 1d), reflecting the partial recovery of microglia towards the end of each ethanol session, as the animals regained consciousness. Microglial cell soma volumes also increased with each ethanol-binge (Fig. 1g, Supp. Fig. 1e, f), which occasionally resulted in drastic shape changes in some cells (Supp. Fig. 2c). Microglial cell density significantly increased on days 5 and 10 (Fig. 1h-i, Supp. Fig. 1g), but some new cells detected on day 5 were absent on day 10, possibly due to cell migration or apoptosis with chronic binge ethanol exposure (Fig. 1h). This progressive density increase correlated with a decrease in microglial surveillance territory (Fig. 1j-m), suggesting a potential adaptative mechanism where microglial proliferation compensates for the loss of ramification and surveillance. Therefore, acute microglial responses to each binge ethanol exposure parallel the onset and recovery from ethanol-induced sedation, and chronic alcohol abuse has a cumulative effect on microglial morphology, distribution, and surveillance territory.

### Macrophage-specific *MyD88* deletion protects from ethanol-induced microglial reactivity, neuronal loss and motor impairments

Next, we investigated the effects of alcohol on neuronal integrity in the same brain region where we observed significant microglial alterations. Immunohistochemical analysis revealed a significant decrease in neuronal nuclei density in the somatosensory cortex of mice subjected to the 10-day ethanol course, compared to saline-treated animals. (Fig. 2a-b). To differentiate between alcohol’s direct neurotoxic effects and indirect effects mediated by inflammation, we genetically targeted *MyD88* in cells of the monocytic lineage, particularly tissue-resident macrophages like Kupffer cells (liver) and microglia (brain). This was achieved using a transgenic mouse line with constitutive macrophage-specific *MyD88* deletion (*Cx3cr1Cre^Tg/0^:MyD88^F/F^*)^57^, which was further crossed to a cre-dependent reporter (*Rosa26-tdTomato^F/F^*) to label all cre^+^ and MyD88-knockout cells. Mice lacking *MyD88* in tissue resident macrophages (Supp. Fig. 3a) were significantly protected from ethanol-induced neuronal cell death even after ten days of alcohol abuse (Fig. 2a-b). Importantly we detected similar protection from neuronal damage in the prefrontal cortex – a brain region critical for executive functions and implicated in alcohol dependence^19^ – which was also comparable to that seen in global *MyD88* knockout mice (Fig. 2c-d). Since global *MyD88* deletion previously protected mice from ethanol-induced motor impairments using the rotarod test^44^, we also assessed motor function in macrophage-specific *MyD88-*knockout mice, using the same assay and ethanol exposure protocol (single dose, 2g/kg BW) as in those earlier studies^44, 58–60^. Genetic ablation of *Myd88* in all macrophages enabled mice to recover from ethanol significantly faster and remain on the rotarod significantly longer than *MyD88^F/F^* controls (Fig. 2e), mirroring the phenotype observed in global knockout animals^44^. These data demonstrate that a macrophage-specific *MyD88*-dependent mechanism is directly involved in ethanol-induced neuronal damage and motor impairments.

**Fig. 2:**
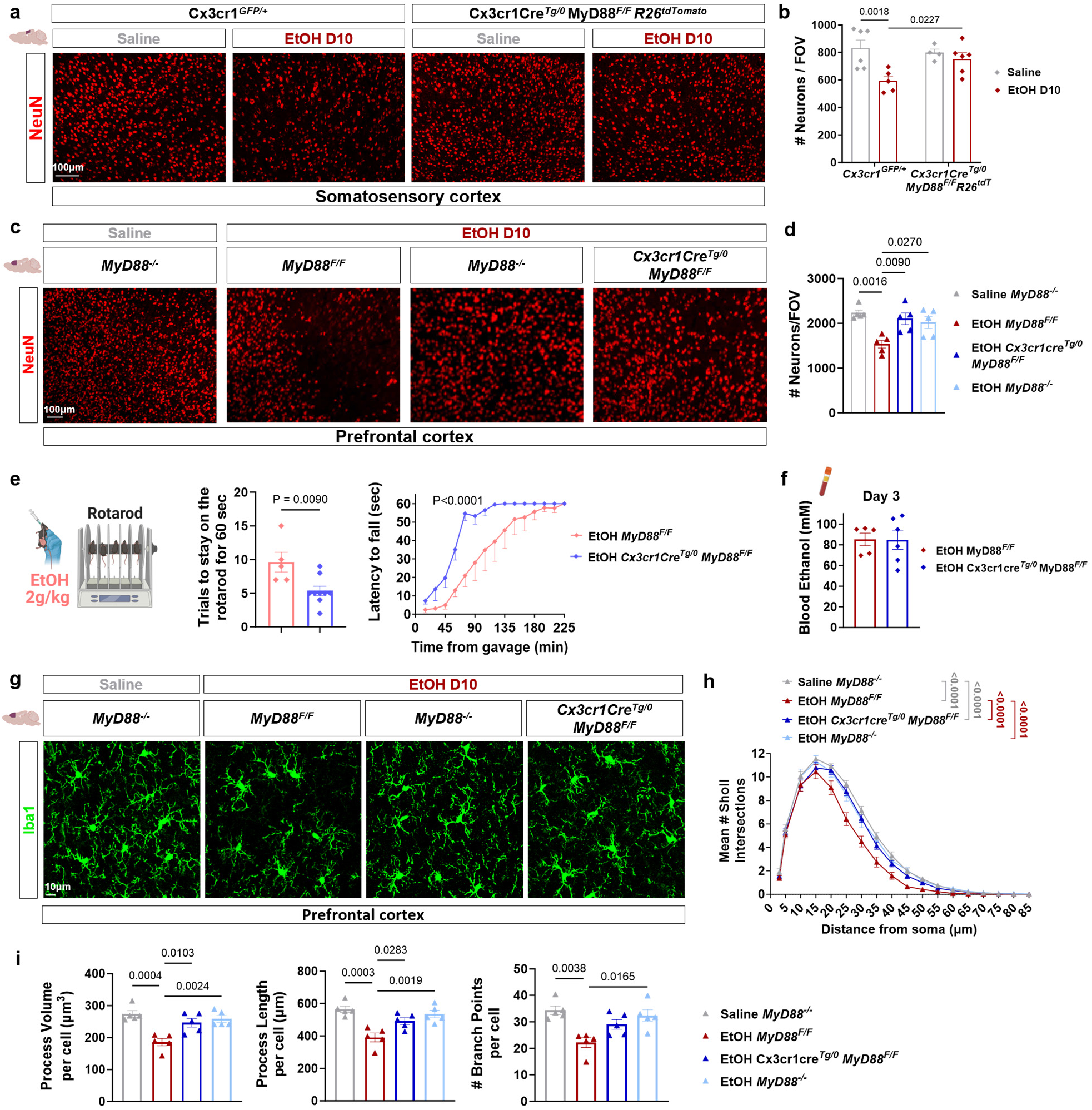
Macrophage-specific *MyD88* deletion protects from ethanol-induced microglial reactivity, neuronal loss and motor impairments. **a.** Representative epifluorescence microscopy images of NeuN-labeled neurons in somatosensory cortex of saline- and ethanol-treated *Cx3cr1^GFP/+^* and *Cx3cr1cre^Tg/0^;MyD88^F/F^;Rosa26^tdTomato^* mice after 10 consecutive days of treatment. Scale bar, 100 µm. **b.** Quantification of NeuN^+^ neurons per field of view (FOV) in somatosensory cortex of mice treated as described in (**a**). n = 4-6 mice per group. Two-way ANOVA with Tukey’s multiple comparison test. **c.** Representative epifluorescence microscopy images of NeuN-labeled neurons in prefrontal cortex of saline- and ethanol-treated *MyD88^-/-^*, *MyD88^F/F^*, and *Cx3cr1cre^Tg/0^;MyD88^F/F^* mice after 10 days of treatment. Scale bar, 100 µm. **d.** Quantification of NeuN^+^ neurons per FOV in prefrontal cortex of mice treated as described in (**c**). n = 5 mice per group. **e.** Rotarod assessment of ethanol-induced motor impairments following a single binge ethanol exposure (2 g/kg BW). Number of trials to stay on the rod and latency to falling off the rotarod for ethanol-treated *MyD88^F/F^* (n=5) and *Cx3cr1cre^Tg/0^;MyD88^F/F^* (n=9) mice. Unpaired t-test and Wilcoxon matched-pairs signed rank test, respectively. **f.** Quantification of blood ethanol levels in plasma samples collected on day 3 of binge ethanol exposure (5 g/kg BW) from *MyD88^F/F^* and *Cx3cr1cre^Tg/0^;MyD88^F/F^* mice. n = 5–6 mice per group. Unpaired t-test. **g.** Representative confocal microscopy images of Iba1-labeled microglia in prefrontal cortex of mice treated as described in (**c**) on day 10 of binge ethanol exposure. Scale bar, 10 µm. **h, i.** Sholl analysis **(h)** and quantification of microglial process volume, process length and branch points **(i)** in prefrontal cortex of mice treated as described in (**c**). n = 5 mice per group. Data presented as mean ± SEM. **d, i:** One-way ANOVA with Tukey’s multiple comparison test; **h:** Two-way ANOVA with Tukey’s multiple comparison test. Only statistically significant comparisons are shown.

Importantly, the MyD88-dependent protection in motor performance was not attributable to altered peripheral ethanol metabolism, as evidenced by comparable liver alcohol dehydrogenase (ADH) activity (Supp. Fig. 3b) and blood ethanol concentration (BEC) between macrophage-specific *MyD88* knockout mice and their genetic controls (Fig. 2f). Prior research showed that MyD88 signaling, particularly in myeloid cells, can drive liver inflammation and damage^61^, which – we reasoned – could indirectly influence neuroinflammation, neuronal health, and animal behavior. We found no significant differences in IL-1β, TNF, or MCP-1 mRNA levels in the liver of ethanol-treated macrophage-specific *MyD88* knockout animals compared to *MyD88^+/+^* controls, after the 10-day alcohol course (Supp. Fig. 3c). Furthermore, plasma levels of the liver damage markers alanine transaminase (ALT) and aspartate transferase (AST) showed no differences (Supp. Fig. 3d), with only a decrease in liver triglycerides (TG) in macrophage-specific knockout animals compared to controls, which was independent of ethanol treatment (Supp. Fig. 3d). Consequently, we reasoned that a brain-specific mechanism was likely responsible for the neuronal protection from alcohol abuse in the macrophage-specific *Myd88* knockout mice.

Notably, histological analysis of microglial morphology in these mice showed significant protection against ethanol-induced reductions in total process coverage, ramification, and cell body size in both the prefrontal and somatosensory cortex, compared to saline-treated controls (Fig. 2g-i; Supp. Fig. 3e-f), which were not inherently altered by *Cx3cr1* or *MyD88* genotype. Consistent with the neuronal survival data, the macrophage-specific *MyD88* deletion conferred comparable protection in microglial morphology metrics as the global knockout. Collectively, these findings suggest that ethanol may impair neuronal survival and function by directly acting on the brain’s primary myeloid-derived immune cell population –microglia– in a MyD88-dependent manner.

### Microglia-specific *MyD88* deletion prevents acute and chronic ethanol-induced microglial responses and neuronal cell death

To study the role of microglial MyD88 in neuroimmune responses to alcohol abuse, we utilized inducible *Cx3cr1cre^ERT2^* mice which specifically target microglia (when used 4-5 weeks after tamoxifen induction, see Methods). To image endogenously labeled *MyD88* knockout microglia *in vivo*, we generated *Cx3cr1cre^ERT2^:Myd88^F/F^:Rosa26-tdTomato^F/F^* mice and evaluated the efficiency of *MyD88* deletion in genomic DNA^62^ from microglia isolated from macrophage- and microglia-specific MyD88-knockout mice (Supp. Fig. 3a). Similar to *Myd88^+/+^* controls (Fig. 1), we used two-photon microscopy to image the somatosensory cortex of awake microglia-specific *MyD88* knockout mice both before and after saline administration. Subsequently, the same brain regions in the same animals were re-imaged on days 1, 3, 5, and 10, both before and after ethanol administration, at the same timepoints post-gavage as the control *Cx3cr1^GFP/+^* mice (Fig. 1a, 3a). During each daily imaging session across the 10-day ethanol course, microglia lacking *MyD88* only showed minor morphological reactivity (Fig. 3b,c), peaking again 1 hour after ethanol administration (Supp. Fig. 4a) and recovering by 5-6 hours post-gavage, while remaining mostly unchanged throughout the saline treatment day (Fig. 3b,c). 3D tracing of microglial processes (Fig. 3d) showed a slight reduction of process volume, length, branch points, and surveillance territory (Fig. 3g) 1 hour after ethanol administration throughout the 10-day course. No differences were observed 5-6 hours post-gavage (Supp. Fig. 4b), reflecting rapid morphological recovery of microglia within each daily session. In contrast to control mice, the microglia-specific *MyD88* knockout animals did not show progressive worsening of microglial reactivity with increasing days of binge ethanol exposure (Fig. 3d,g). Microglial cell density and mean cell body volume also remained unchanged throughout the 10-day ethanol course (Fig. 3f, g, Supp. Fig. 4). Compared to controls, microglia-specific *MyD88* knockout mice exhibited significant attenuation of all ethanol-induced microglial morphological activation metrics (Fig. 3g). Importantly, deleting *Myd88* only from microglia was also sufficient to protect from the extensive neuronal cell death that control animals experienced after 10 consecutive binge doses of ethanol (5g/kg BW, Fig. 3h-i). These results demonstrate that microglial *MyD88* is required for both acute and chronic microglial responses to excessive ethanol exposure, and for mediating neuronal damage in the prefrontal cortex following chronic binge drinking.

**Fig. 3:**
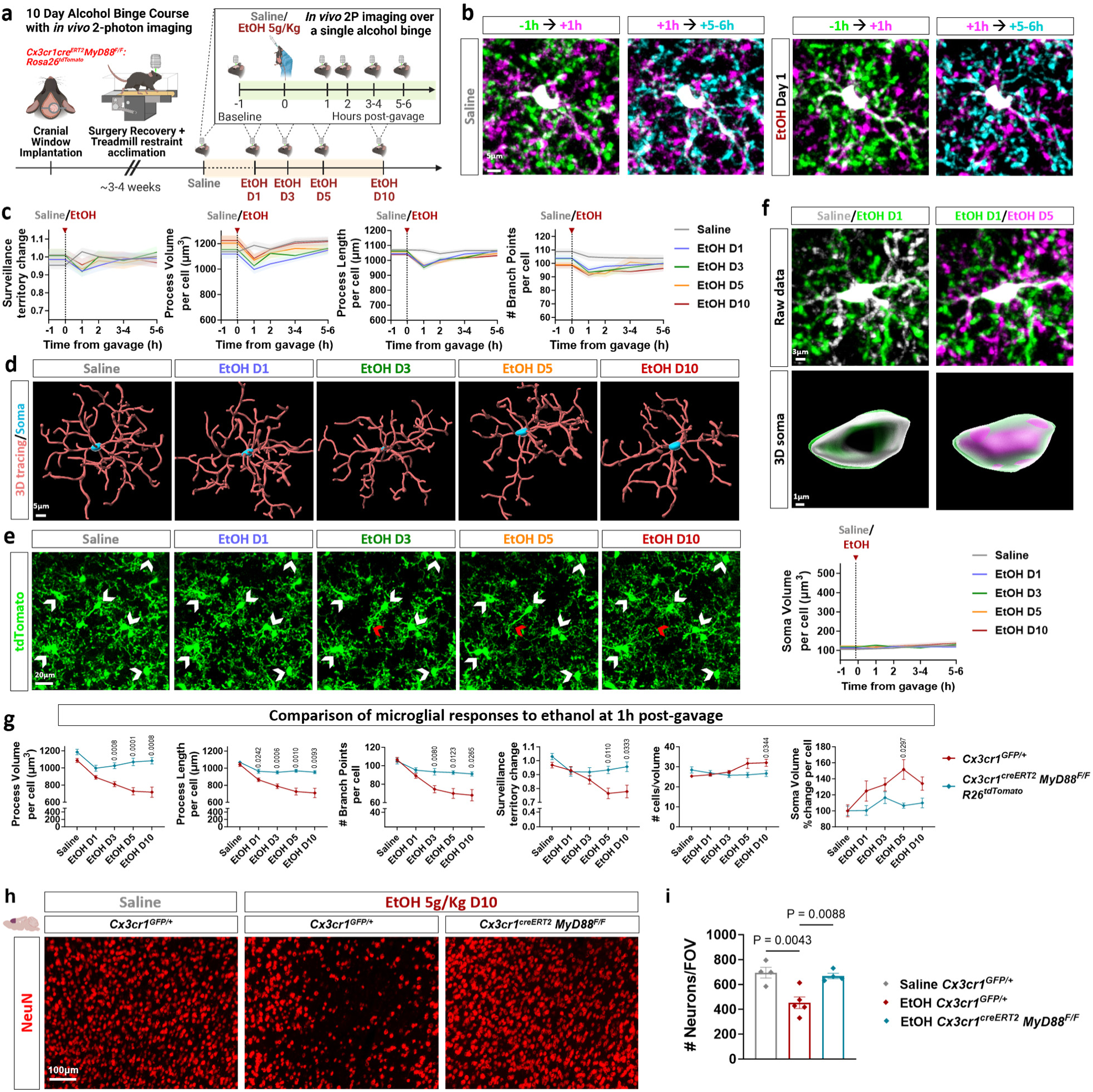
Microglia-specific *MyD88* deletion prevents acute and chronic ethanol-induced microglial responses *in vivo* and neuronal cell death. **a.** Schematic representation of *in vivo* two-photon microscopy experimental design for imaging the evolution of microglial responses in somatosensory cortex of the same *Cx3cr1^creERT2^;MyD88^F/F^;Rosa26^tdTomato^* mice, before (saline) and during acute and chronic binge ethanol exposure (5 g/kg BW). **b.** Representative two-photon microscopy images of the same microglial cell, captured at -1 h, +1 h and +5–6 h following gavage on saline treatment day and on ethanol day 1. Scale bar, 5 µm. **c.** Time course of changes in microglial surveillance territory, process volume, process length, and number of branch points following gavage on saline day and on binge ethanol exposure days 1, 3, 5, and 10. **d.** Representative 3D tracings of microglial processes and cell soma reconstructions from the same cell at 1h post-gavage, on saline day and on ethanol exposure days 1, 3, 5, and 10. Scale bar, 5µm. **e.** Representative two-photon microscopy images of microglia in the same cortical volume at 1h post-gavage, on saline treatment day and on binge ethanol exposure days 1, 3, 5, and 10. White arrowheads indicate cells tracked throughout all days of imaging. Red arrowheads mark the location of a cell that is absent after ethanol day 3. Scale bar, 20 µm. **f.** Representative two-photon microscopy images of the same microglial cell and its 3D reconstructed cell soma at 1h post-gavage on saline treatment day and binge ethanol exposure days 1 and 5. Scale bars, 3 µm (raw data), 1 µm (3D soma). Graph shows the quantification of cell soma volume over hours following gavage on saline day and on ethanol exposure days 1, 3, 5, and 10. **g.** Comparison of microglial process volume, process length, branch points, surveillance territory change, cell density and soma volume percent change at 1h post-gavage on saline day and on binge ethanol exposure days 1, 3, 5, and 10 between *Cx3cr1^GFP/+^* and *Cx3cr1^creERT2^;MyD88^F/F^;Rosa26^tdTomato^* mice. Mixed effects analysis with Sidak’s multiple comparison test. **h.** Representative epifluorescence microscopy images of NeuN-labeled neurons in prefrontal cortex of saline- and ethanol-exposed (5 g/kg BW) mice on day 10 of treatment. Scale bar, 100 µm. **i.** Quantification of NeuN^+^ neurons per field of view (FOV) in prefrontal cortex of mice treated as described in (**h**). n = 4-5 mice per group. One-way ANOVA with Tukey’s multiple comparison test. Data presented as mean ± SEM. **c, f-g:** n = 11 imaged locations from 6 mice per group. Only statistically significant comparisons are shown.

### Microglial *MyD88* deletion protects from chronic ethanol-induced synapse elimination

We next explored whether microglia also respond to lower, non-sedative amounts of ethanol, and whether such responses could still impact the neuronal network in the absence of a neurotoxic ethanol challenge. We used the same 10-day chronic exposure model, but administered 2g ethanol / kg BW, a dose sufficient to cause motor impairments (Fig. 2e), as also shown previously^44, 58–60^. This dose did not cause neuronal loss in the prefrontal cortex even after 10 days of treatment, in WT or in either of the two microglia-specific *MyD88* knockout mouse lines (Supp. Fig. 5a, b). We next hypothesized that even though the lower dose of ethanol does not kill neurons, it might still impact their function at the synaptic level. We thus performed immunofluorescence followed by high-resolution confocal microscopy for the pre- and post-synaptic proteins synaptophysin (Syn) and Homer1, respectively. Ethanol-treated mice showed a reduction in the number of Homer1^+^:Syn^+^ colocalized puncta (indicating synapses), and in Homer1^+^ and Syn^+^ volumes compared to saline controls, all of which were significantly restored in both microglia-specific MyD88 knockout lines (Fig. 4a, b, Supp. Fig. 5c).

**Fig. 4:**
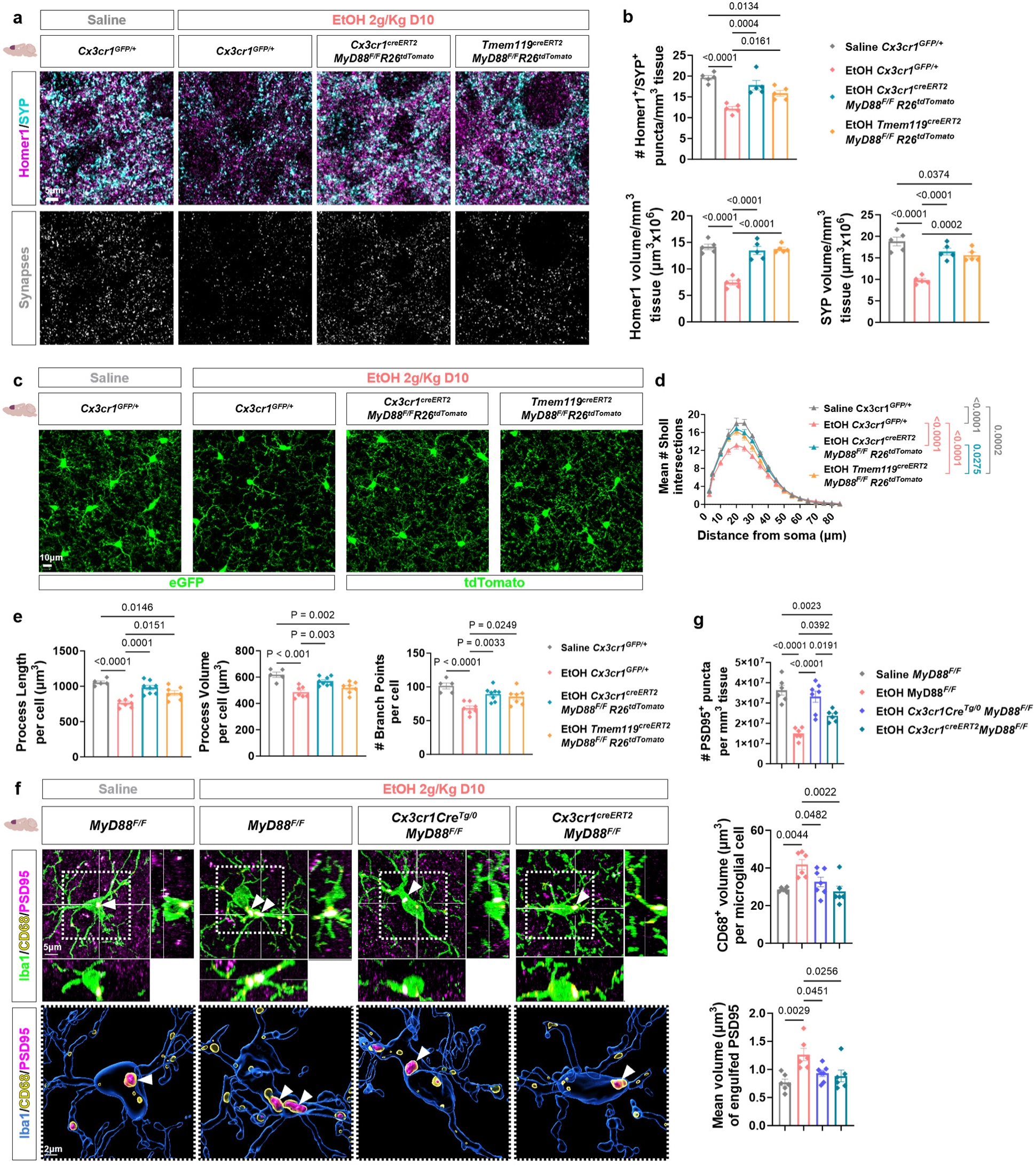
Microglial *MyD88* deletion protects from chronic ethanol-induced synapse elimination. **a.** Representative confocal microscopy images of post-synaptic Homer1 and pre-synaptic synaptophysin (SYP) in prefrontal cortex of mice orally administered saline or ethanol (2g/kg BW) for 10 days. Colocalized double-positive puncta represent synapses. Scale bar, 5 µm. **b.** Volumetric analysis of Homer1 and synaptophysin immunofluorescence, and quantification of colocalized puncta in prefrontal cortex of mice treated as described in (**a**). n= 5 mice per group. **c.** Representative confocal microscopy images of eGFP- and tdTomato-labeled microglia in prefrontal cortex of mice treated as described in (**a**). Scale bar, 10 µm. **d-e.** Sholl analysis **(d)** and quantification of microglial process length, process volume and branch points **(e)** of microglia in prefrontal cortex of mice treated as described in (**a**). n= 5-8 mice per group. **f.** Representative confocal microscopy images (top) of Iba1^+^ microglia, CD68^+^ lysosomes and PSD-95^+^ post-synaptic densities in the prefrontal cortex of mice treated as described in (**a**). Bottom panels show 3D reconstructions highlighting the engulfed synaptic material within microglial lysosomes shown in the boxed regions of the images above them. Scale bars, 5 µm (top row) and 2 µm (bottom row). **g**. Quantification of PSD-95^+^ puncta per tissue volume, CD68^+^ lysosome volume per Iba1^+^ microglial cell, and mean volume of phagocytosed PSD-95 (PSD-95^+^ particles within CD68^+^ microglial lysosomes), in prefrontal cortex of mice treated as described in (**a**). n = 6-7 mice per group. Data presented as mean ± SEM; **b, e, g:** One-way ANOVA with Tukey’s multiple comparison test. **d:** Two-way ANOVA with Tukey’s multiple comparison test. Only statistically significant comparisons are shown.

Interestingly, even this lower dose of ethanol elicited a milder but significant microglial response in the prefrontal cortex. Compared to saline-treated controls, ethanol-treated *Cx3cr1^GFP/+^* mice showed loss of ramification (Fig. 4c) and reductions in Sholl intersections, process length, volume, and number of branch points (Fig. 4d, e). Conditional deletion of *MyD88* using either microglia-specific cre recombinase line significantly restored their morphology to almost saline levels, despite ethanol exposure for 10 days (Fig. 4d, e).

Alcohol intake has been previously reported to induce aberrant synaptic pruning by microglia in the prefrontal cortex in a TNF-dependent manner^63^, and in the hippocampus in a TREM2-dependent manner^64^. We thus hypothesized that the ethanol-induced *MyD88*-dependent synapse loss we observed in the prefrontal cortex could be the result of increased removal of synaptic structures by microglia. We used immunofluorescence labeling of post-synaptic density-95 (PSD-95), CD68^+^ phagolysosomes, and Iba1^+^ microglia on brain tissue sections from saline- and ethanol-treated (2g/kg BW) *MyD88^F/F^* and conditional knockout mice (Fig. 4f). The number of PSD-95 puncta in the prefrontal cortex was significantly decreased by ethanol treatment, and both macrophage- and microglia-specific deletion of *MyD88* significantly restored the loss of PSD-95 puncta (Fig. 4g, Supp. Fig. 5d). Volumetric analysis of CD68 staining revealed an ethanol-induced increase in CD68^+^ volume per Iba1^+^ microglial cell, which was prevented by macrophage- and microglia-specific *MyD88* deletion (Fig. 4g, Supp. Fig. 5d). Importantly, the mean volume of PSD-95^+^ puncta detected within CD68^+^ phagolysosomes in Iba1^+^ microglia was significantly increased in ethanol-treated *MyD88^F/F^* mice but restored in both conditional MyD88-knockout animals (Fig. 4g). These findings indicate that microglial signaling through MyD88 mediates synaptic loss in the prefrontal cortex following chronic alcohol consumption.

### Ethanol-induced loss of neuronal activity is dependent on microglial MyD88

In addition to their role in synapse remodeling, microglia can also respond to and modulate neuronal activity^31, 35, 37, 38^. We thus explored whether microglia might impact neuronal function in the context of acute and chronic alcohol abuse, by performing longitudinal *in vivo* calcium imaging in the cortex of control (*MyD88^+/+^*) and microglia-specific *MyD88* knockout mice (Fig. 5a). One week after cranial window implantation over the somatosensory cortex, all animals were intravenously injected with adeno-associated virus (AAV) expressing CaMPARI (Calcium-Modulated Photoactivatable Ratiometric Integrator), a genetically encoded calcium indicator that selectively labels active neurons^65, 66^. Upon exposure to 400 nm light in a high-calcium environment, CaMPARI undergoes irreversible photoconversion from green to red fluorescence. This process marks neurons actively firing action potentials during the illumination period, allowing for time-stamped detection of neuronal activity across large brain volumes in freely moving animals without requiring simultaneous imaging^67^. To leverage this, each photoconversion session in our study involved a 10 min illumination of 400 nm light, delivered 1-2 hours post-gavage to awake, head-fixed animals on a treadmill (Fig. 5a). Subsequently, *in vivo* two-photon microscopy was used to record the green and red fluorescence from neurons across multiple cortical planes, spanning the somatosensory cortex of each mouse. Furthermore, the significant decay of the converted red signal (approximately 95% within one week^67^) allowed us to perform multiple recording sessions and re-evaluate neuronal activity in the same brain regions of the same mice: first after saline administration, then 10 days later – after the initial ethanol treatment– and finally after the last ethanol dose on day 10 of the binge drinking course. After image acquisition, a total of 43,029 neuronal cell bodies were segmented from all images acquired from wild-type and *Cx3cr1^creERT2^:MyD88^F/F^* mice (Fig. 5a). Although the number of segmented cells per imaged volume varied, their distributions were similar between wild-type and microglia-specific *MyD88* knockout mice under both saline and ethanol conditions (Fig. 5b), allowing for a fair comparison between the cohorts. In most cases, the same cells in the same cortical locations were successfully identified and reimaged across all three sessions (Fig. 5c) allowing for assessment of neuronal activity changes from the same neurons on saline and ethanol days 1 and 10. Comparison of cellular red to green (RG) ratios across conditions revealed that *MyD88^+/+^* mice showed significantly reduced activity following both 1 and 10 days of ethanol treatment, compared to their own saline recordings (Fig. 5d, e). In contrast, microglia-specific *MyD88* knockout mice maintained consistent overall neuronal activity between saline and ethanol day 1 and in some cases even showed an increase on ethanol day 10, indicating resistance to ethanol-induced neuronal activity decline (Fig. 5d, e). These results suggest that microglial *MyD88* is required for both acute and chronic ethanol-induced reductions in neuronal activity.

**Fig. 5:**
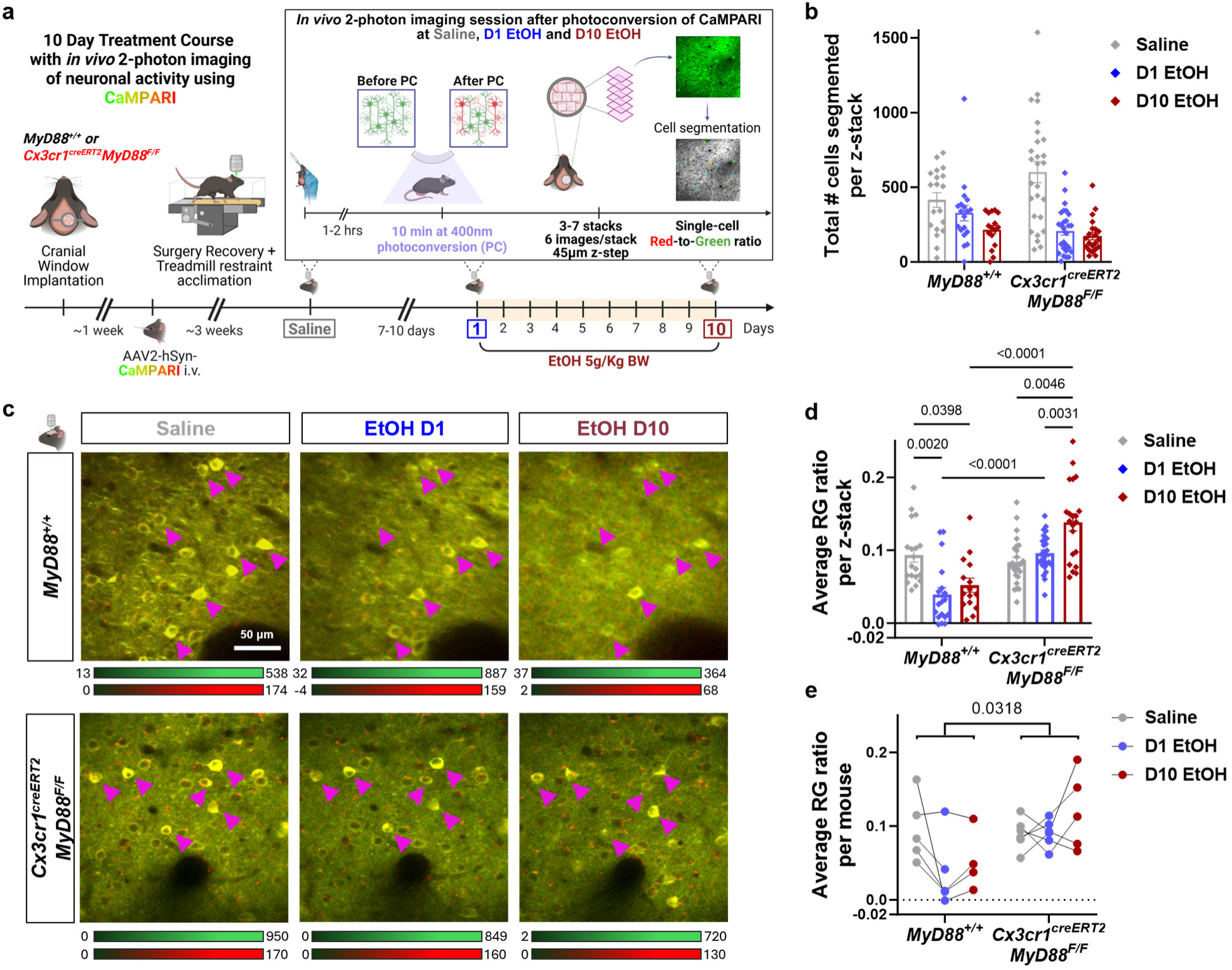
Ethanol-induced loss of neuronal activity is dependent on microglial MyD88. **a.** Experimental setup for CaMPARI photoconversion and *in vivo* two-photon microscopy recording of neuronal activity. Mice underwent implantation of cranial windows over somatosensory cortex, followed by administration of AAV expressing the Ca^+2^ indicator CaMPARI. After recovery, mice underwent 3x 10-min cycles of CaMPARI photoconversion 9-10 days apart, first following a single dose of saline and then on the 1^st^ and 10^th^ day of ethanol treatment (5g/kg BW), all while on a treadmill. After photoconversion, the mice were imaged *in vivo*, under a two-photon microscope. 3-7 cortical columns (z-stacks, each containing 6 z-planes, 45 μm apart) were imaged and the red-to-green (RG) ratio of individually segmented neuronal cell bodies was calculated. **b.** Total number of neurons segmented per analyzed column in *WT* and *Cx3cr1^creERT2^;MyD88^F/F^* mice treated as described in (**a**). **c.** Representative *in vivo* two-photon microscopy images of CaMPARI photoconverted cells in mice treated as described in (**a**). The same cortical locations and photoconverted cells (arrows) are shown for each group across treatments. Scale bar, 50 µm. **d.** Average RG ratios per analyzed column for each genotype/treatment group. **e.** Average RG ratio changes per animal across saline and ethanol exposure days, highlighting significant differences between genotypes. Data presented as mean ± SEM. **b, d-e:** n = 15-19 z-stacks from 4-5 WT mice, n = 21-27 z-stacks from 5-6 Cx3cr1^creERT2^;MyD88^F/F^ mice across conditions. Mixed-effects model with Tukey’s multiple comparison test. Statistically significant comparisons are shown.

### Microglia-specific *MyD88* deletion prevents ethanol-induced motor impairments and sedation

To investigate whether the observed neuronal activity changes influenced alcohol-related behaviors, we evaluated motor performance (rotarod), locomotor activity, and ethanol-induced sedation (Fig. 6a) in control and microglia-specific *MyD88* knockout mice. In the rotarod test, *Cx3cr1cre^ERT2^:MyD88^F/F^* mice required significantly fewer trials to stay on the rotarod for 60 seconds and exhibited increased latency to falling off the rotarod compared to *Cx3cr1^GFP/+^* ethanol-treated controls (Fig. 6b). The *Tmem119cre^ERT2^:MyD88^F/F^* mice treated with ethanol were less protected from ethanol-induced motor impairments in this test, but still showed significantly increased latency to falling off the rotarod compared to *Cx3cr1^GFP/+^* mice (Fig. 6b). In the locomotor activity test, both microglia-specific *MyD88* knockout lines showed a significant increase in total beam breaks indicative of locomotor activity throughout the two hours following 2g/kg BW ethanol administration compared to controls (Fig. 6c). Interestingly, *Tmem119cre^ERT2^:MyD88^F/F^* mice with improved microglial specificity were even less affected by ethanol than the *Cx3cr1cre^ERT2^:MyD88^F/F^* mice showing improved locomotor activity (Fig. 6c). To test their resistance from alcohol-induced impairments further, we acutely injected another cohort of *Tmem119cre^ERT2^:MyD88^F/F^* mice with a sedative dose of ethanol (3.5g/kg BW) intraperitoneally and performed the loss of righting reflex (LORR) test, commonly used to evaluate ethanol-induced sedation^44, 60^. Microglia-specific *MyD88* knockout mice showed significantly increased latency to LORR (Fig. 6d) and recovered their righting reflex in almost half the time of *MyD88^F/F^* control mice, demonstrating significant protection even from alcohol’s sedative effects (Fig. 6d).

**Fig. 6:**
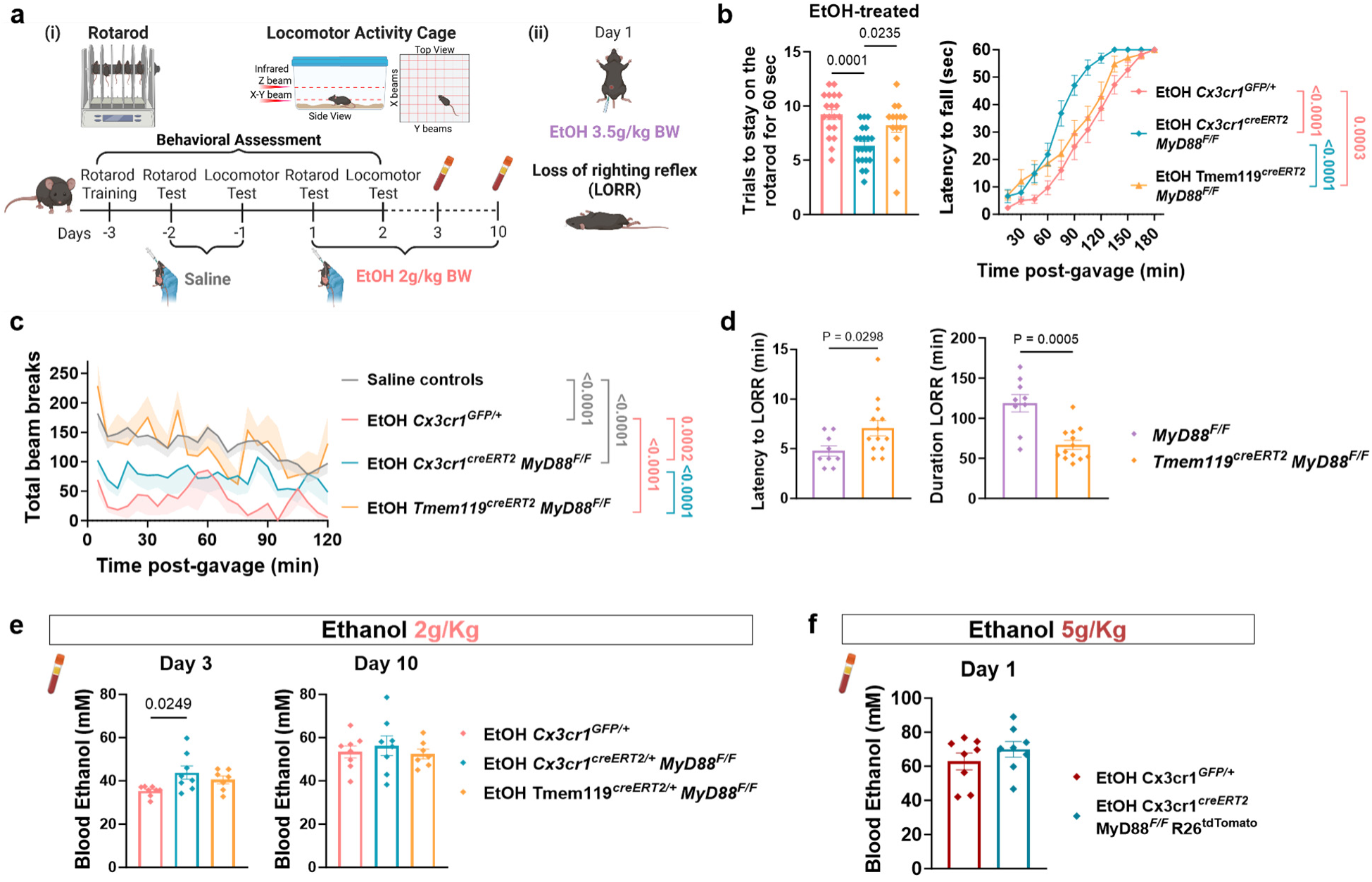
Microglia-specific *MyD88* deletion prevents ethanol-induced motor impairments and sedation. **a.** Schematic diagram of experimental design for the assessment of ethanol-induced motor impairments. **(i)** rotarod and locomotor activity cage assessments performed after the first and second day of binge ethanol oral administration (2 g/kg BW). **(ii)** Loss of righting reflex (LORR) assessment after a single intraperitoneal injection of 3.5 g EtOH/kg BW. **b.** Number of trials to stay on the rod and latency to falling off the rotarod on day 1 of the 10-day ethanol course. n = 19 ethanol-treated *Cx3cr1^GFP/+^*, n = 21 *Cx3cr1^creERT2^;MyD88^F/F^* and n = 14 *Tmem119^creERT2^;MyD88^F/F^* mice. One-way ANOVA with Tukey’s multiple comparison test (trials to stay), and Friedman with Dunn’s multiple comparisons test (latency to fall). **c.** Total beam breaks recorded in the locomotor activity cage over 2 hours after ethanol administration on day 2 of the 10-day ethanol course. Saline control data were collected one day before ethanol course initiation, as described in (**a**(**i**)). Data binned every 5 minutes. n = 5 *Cx3cr1^GFP/+^*, n = 9 *Cx3cr1^creERT2^;MyD88^F/F^*, n = 8 *Tmem119^creERT2^;MyD88^F/F^* mice. Kruskal-Wallis with Dunn’s multiple comparisons test. **d.** Latency to LORR and duration of LORR following treatment as described in (**a(ii)**). n = 9 *MyD88^F/F^* and n = 13 *Tmem119^creERT2^;MyD88^F/F^* mice. Unpaired t-test. **e.** Quantification of blood ethanol concentration in plasma samples on day 3 and 10 of binge ethanol exposure (2g/kg BW, n = 7-8 mice / group). One-way ANOVA with Tukey’s multiple comparison test. **f.** Quantification of blood ethanol concentration in plasma samples on day 1 of binge ethanol exposure (5g/kg BW). n = 8 mice/group. Unpaired t-test. Data presented as mean ± SEM. Only statistically significant comparisons are shown.

To examine whether differences in peripheral ethanol metabolism or ethanol-induced liver damage could contribute to the observed protection against ethanol-induced behavioral impairments in microglia-specific *MyD88* knockout mice, we measured the levels of ADH and TG in the liver and ALT and AST in the plasma of mice treated with 2g/kg BW ethanol dose. Microglia-specific *MyD88* deletion did not alter liver TG or plasma ALT or AST levels in either conditional knockout line compared to ethanol-treated controls (Supp. Fig. 6a), indicating no evident differences in liver damage between these cohorts. Notably, the levels of liver TG were similarly increased due to ethanol treatment among all groups compared to saline treatment (Supp. Fig. 6a). Liver ADH also showed no difference between control and either conditional *MyD88* knockout cohort (Supp. Fig. 6a). Crucially, BEC levels assessed at 90 minutes post-ethanol administration were consistently similar (or slightly higher) in microglia-specific *MyD88* knockout lines compared to controls across various ethanol exposure paradigms (single, 3, or 10 consecutive binges, and different doses, Fig. 6e, f). Therefore, alcohol abuse causes motor and cognitive impairments in a microglial-dependent manner, and disrupting MyD88-mediated signaling selectively in microglia protects against ethanol-induced intoxication without affecting liver damage or ethanol metabolism.

### Alcohol abuse induces a phagocytosis, matrisome, and cell growth/adhesion related transcriptional signature in microglia

To better understand how microglia could exert such profound changes in neuronal structure and function following alcohol abuse, we performed bulk RNA sequencing on microglia isolated from the whole brain of saline- and ethanol-treated mice. We collected microglia on day 5, 18 hours after the fourth daily ethanol binge (Fig. 7a), a time point chosen because (i) our *in vivo* imaging revealed peak microglial reactivity on day 5 of the 10-day binge model (Fig. 1), and (ii) previous studies showed brain inflammatory gene expression peaks 18 hours after acute binge ethanol administration^68^. Differential expression (DE) analysis identified 91 genes with significantly altered expression (padj<0.05, |log2FC|>0.6) between saline- and ethanol-treated mice (Fig. 7b). Binge ethanol exposure significantly downregulated genes associated with the immune response (*Tnfsf12*, *Ccr1*, *Ccr1l1*) and transcriptional regulation (*Klf13*, *Hlf*, *Jade2*, *Kdm6b*), while it upregulated genes involved in heme metabolism/mitochondrial dynamics (*Hebp1, Aldh1l2*), phagocytosis (*Cd36, Eps8*), extracellular matrix (ECM) regulation (*Dcn,Plxnd1*), DNA replication (*Mcm2*, *Mcm5*) and cytoskeletal dynamics/cell adhesion (*Septin9, Tiam1, Pdgfrb*). Moreover, pathway analysis showed enrichment in ethanol-exposed microglia for pathways related to phagocytosis, ECM binding and structure, cell growth and adhesion, and apoptosis (Fig. 7c-f). The upregulation of cell growth pathways could explain the microglial cell expansion observed in ethanol-treated mice *in vivo* (Fig. 1h, i), while the upregulation of phagosome/lysosomal pathways supports our histological data showing increased microglial CD68^+^ phagolysosomes containing synaptic material (Fig. 4f, g). Only a limited number of pathways were significantly downregulated, including TNF signaling and circadian rhythm regulation (Fig. 7e-f).

**Fig. 7:**
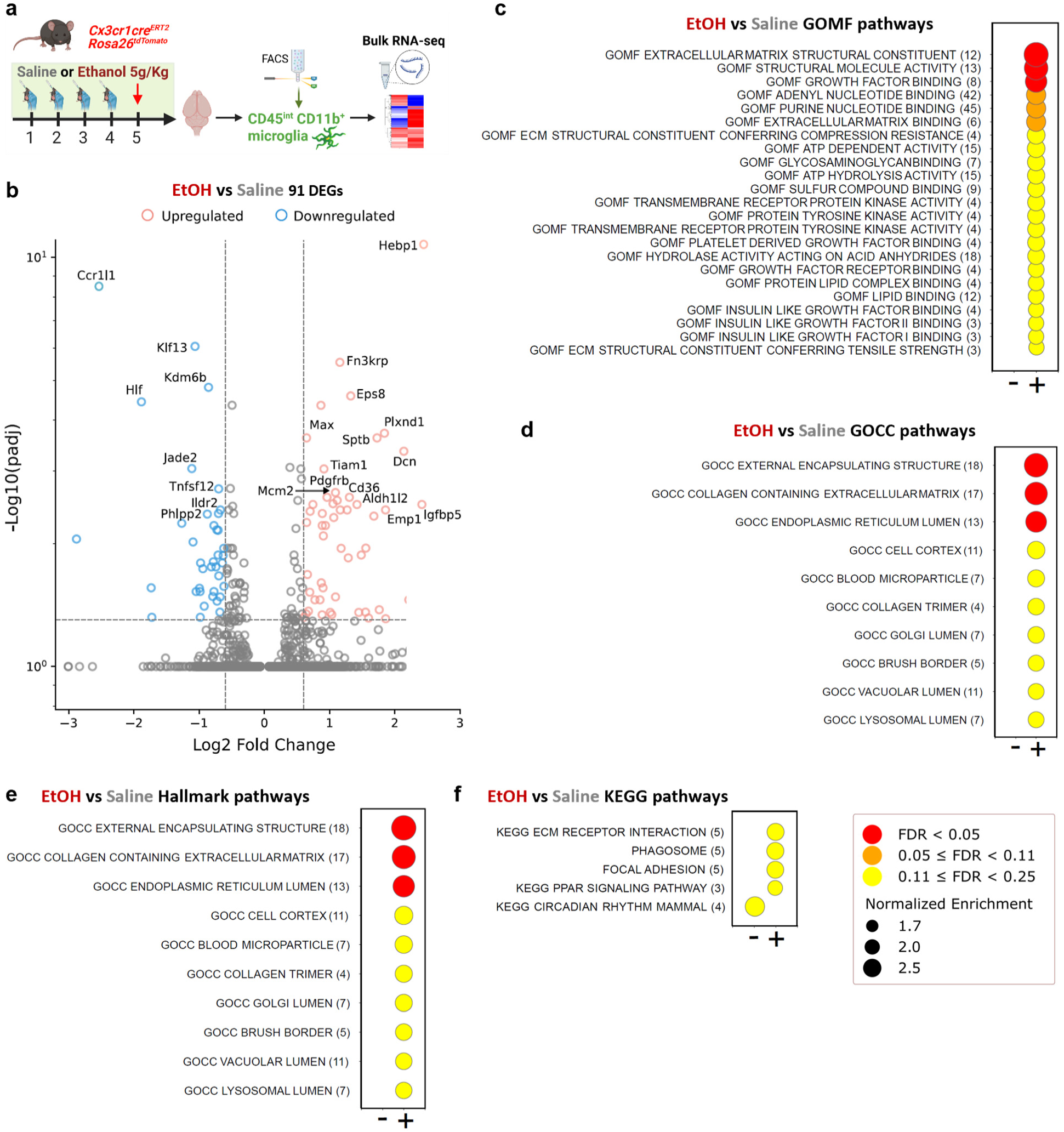
Microglia exhibit a phagocytosis, ECM, and cell growth/adhesion related gene signature following ethanol abuse. **a.** Experimental design for RNAseq experiment in FACS-sorted microglia from *Cx3cr1^creERT2^;Rosa26^tdTomato^* mice administered saline or ethanol (5g/kg BW) daily, with tissue collection 18h after ethanol gavage on day 4 of the ethanol course. **b.** Volcano plot showing significantly differentially expressed genes (DEGs) in ethanol- versus saline-treated mice. Significance thresholds: padj < 0.05, |0.6| < log2FC. n = 5 ethanol day 5 mice and 6 saline day 5-10 mice. **c-f.** Dot plots of significantly positively (+) and negatively (-) enriched pathways from different pathway databases in ethanol-treated mice compared to saline-treated controls.

## Discussion

Alcohol represents the fourth leading preventable cause of death in the United States. Of greatest concern is binge drinking, which impacts 60.4 million adults ages 18 and older, with young adults ages 18-25 showing particularly alarming binge drinking rates of 28.7%^1^. Equally troubling are emerging trends among older adults (12.0% of those 65+ reporting binge drinking) who face heightened risks due to medication interactions, and among women, with approximately one in four female drinkers reporting binge episodes^1^. Research demonstrates that people consuming alcohol at twice the binge threshold are 70 times more likely to have alcohol-related emergency department visits, while those at three times the threshold are 93 times more likely. High-intensity drinking is most common among young adults attending college but affects approximately 40% of all binge drinkers at least occasionally, substantially increasing risks of severe health and safety consequences^1, 2, 69^.

To model these distinct drinking patterns, our 10-day gavage protocol employs either 2 or 5g/kg BW ethanol doses, with the lower dose mimicking standard binge episodes and the higher dose modeling high-intensity drinking that generates BACs exceeding 0.30% (approximately 65mM) ^70, 71^. This approach leverages the accelerated ethanol metabolism of mice —up to 5 times faster than humans^70, 72^ —allowing us to study cumulative effects of repeated intoxication cycles in a compressed timeframe. Despite pharmacokinetic differences, the neurobiological mechanisms affected by ethanol are remarkably conserved across species^73^. Furthermore, the controlled nature of gavage administration ensures precise dosing and consistent BAC profiles, unlike voluntary drinking models where consumption patterns vary, enabling us to explore dose-dependent relationships between exposure intensity and both cell type- and tissue-specific effects. By comparing these two dosing levels, we could distinguish between effects occurring at typical binge levels versus high-intensity drinking, providing translational insights into neurobiological mechanisms that may underlie the detrimental brain impact of alcohol abuse, which has been increasing in prevalence over recent years. We particularly focused on alcohol-induced neuroinflammation, which has emerged as a crucial component of alcohol-related brain pathology^7^, yet the mechanistic links between alcohol abuse, microglial activation, and neuronal dysfunction remain elusive. Our findings establish microglia as central mediators of alcohol’s detrimental neurological effects through a MyD88-dependent mechanism that destabilizes synaptic connections, eventually leading to neuronal loss. This work provides several conceptual advances in understanding how alcohol abuse impacts brain function, both acutely and chronically.

First, our longitudinal *in vivo* imaging recordings reveal that microglial responses to ethanol are temporally coupled to the onset, duration, and resolution of alcohol’s sedative effects. Previous studies documented microglial morphological changes after alcohol exposure^10, 45–48, 54, 74, 75^, but did not connect these changes to behavioral states. We found microglial ramification decreasing within approximately one hour of alcohol administration, preceding peak sedation by 1-2h, and gradually recovering as animals regained consciousness. This dynamic relationship suggests microglia are not merely responding to alcohol-induced brain damage but could be real-time participants in the acute manifestation of alcohol’s neurological effects. This challenges the conventional view that alcohol’s acute effects are solely mediated through direct modulation of neuronal GABA and NMDA receptors^11, 76^. Furthermore, the progressive nature of microglial morphological reactivity across repeated alcohol binge exposure provides crucial insight into how chronic alcohol abuse might drive cumulative brain dysfunction. While reduced microglial motility and surveillance in the mouse visual cortex were previously reported at 1-hour following a single alcohol binge^41^, our data demonstrate that chronic binge exposure leads to adaptive changes in microglial distribution and surveillance patterns. The increased microglial density despite reduced per-cell surveillance capacity suggests a compensatory response that maintains tissue monitoring despite progressively impaired individual cell function.

Previous studies demonstrated that global deletion of TLR4 or MyD88 protected against alcohol-induced brain inflammatory responses and behavioral impairments^10, 43, 44, 77^, but could not distinguish direct effects on neurons from effects mediated by inflammatory cells. Our targeted genetic approach demonstrates that microglia are critical for the deleterious impact of alcohol abuse on brain function, as microglia-specific *MyD88* deletion prevented both morphological microglial activation and loss of neuronal structure and function. This was pronounced in both the somatosensory and the prefrontal cortices, brain regions with increased vulnerability to alcohol in humans^19–22^. These findings align with recent transcriptomic analyses showing upregulation of TLR pathways in prefrontal cortical microglia after alcohol exposure^49, 50^, but also provide causal demonstration that activation of MyD88-dependent pathways in microglia is required for alcohol’s neurological effects.

Alcohol abuse was previously shown to increase microglial engulfment of synaptic structures in the prefrontal cortex^63^ and hippocampus^64^ through TNF or TREM2-dependent mechanisms, respectively. We also showed prominent microglia-mediated synapse engulfment in the prefrontal cortex after 10 days of repeated binge ethanol exposure that is dependent on microglial MyD88. Furthermore, our demonstration that microglial MyD88 deletion protected against ethanol-induced reductions in neuronal activity establishes microglia as regulators of not only structural but also functional neuronal properties in the context of alcohol abuse. Previous studies showed that alcohol decreases excitability in the prefrontal cortex of humans^78^ and animal models^79^, but attributed this primarily to direct effects on neuronal receptors. Our finding that preventing microglial activation preserves neuronal activity despite alcohol abuse indicates that microglial MyD88 signaling significantly contributes to alcohol-induced neuronal silencing. This expands the known roles of microglia in modulating neuronal activity, which have been described in various conditions including general anesthesia^35^, epilepsy^17^, and homeostatic regulation^25, 26^.

Interestingly, we found that ethanol-exposed microglia featured upregulation of phagocytic and ECM-related genes and pathways. These data led us to speculate that microglia may translate heavy alcohol exposure into neuronal dysfunction by facilitating ethanol-induced synaptic elimination and ECM remodeling. This is notable given emerging evidence that the ECM plays critical roles in synapse stabilization^80, 81^, and neuronal function^82, 83^. Furthermore, it was recently shown that microglia can regulate synaptic plasticity through ECM remodeling^84^. Ethanol exposure induces degradation of interstitial matrix and basement membrane structural proteins through upregulation of proteolytic enzyme activity^85^. For example, increased matrix metalloproteinase-9 has been previously reported in the prefrontal cortex of people with a history of alcohol abuse^86, 87^, suggesting that ECM degradation is a conserved pathological consequence of alcohol abuse across species. Paradoxically ethanol exposure increased perineuronal net components (specialized ECM assemblies ensheathing inhibitory neurons) in the insular cortex^88^, suggesting that alcohol’s effects on the brain ECM likely involve complex regulatory mechanisms and require further investigation. Human microglia derived from induced pluripotent stem cells of individuals with AUD also show enhanced expression of phagocytosis- and MHCII complex-related genes and promote synapse elimination in co-culture with neurons^89^, in agreement with our data. In addition, our findings have significant implications for therapeutic approaches to AUD. The identification of microglial MyD88 signaling as a critical mediator of alcohol abuse effects suggests that targeting this pathway might mitigate both acute intoxication and chronic neurodegeneration. Importantly, this approach would differ fundamentally from existing pharmacotherapies for AUD, which primarily target neuronal receptors or reward circuitry^90, 91^.

Several questions remain for future investigation. First, the upstream signals that activate microglial MyD88 after alcohol abuse merit closer examination. While TLR4 has been the most studied receptor in the context of AUD^10^, other pattern recognition receptors utilizing MyD88, including TLR2, may also contribute. Indeed, alcohol induces TLR4 and TLR2 recruitment into lipid rafts in cultured microglia and promotes their physical interaction, which coincides with inflammatory mediator release^92^. Second, determining whether the observed microglial responses represent pathological activation or attempted repair, and the precise temporal relationship between microglial responses, synaptic elimination, and neuronal silencing across different brain regions, ages, and sexes warrant further exploration. Third, investigating whether microglial MyD88 is involved in addiction, withdrawal, or alcohol-seeking behaviors will be crucial for guiding future therapeutic strategies, given that MyD88 induction correlates with lifetime alcohol consumption, age of drinking onset, and neurodegeneration in people with AUD^93^.

In conclusion, our study establishes microglia as central mediators of alcohol’s neurological effects through MyD88-dependent synaptic dysfunction and neuronal damage. These findings demonstrate that modulating microglial responses to alcohol abuse might address the underlying neuroimmune component of AUD and protect against neuronal structural and functional disruptions while preserving beneficial microglial functions. Our work reveals a novel mechanistic framework for understanding alcohol neuropathology that integrates immune signaling, neuronal integrity and function, and animal behavior, opening new avenues for therapeutic intervention targeting the neuroimmune response in alcohol use disorder.

## Data availability

All data that support the findings of this study will be available after publication from the corresponding author upon reasonable request.

## Acknowledgements

We thank Catherine Richards and Emma Wozencraft for animal breeding, and PCR experiments; Dr. Thomas Jaramillo, (Rodent Behavior Core Cleveland Clinic) for behavioral experiment support; the Flow Cytometry and Genomics Cores at Cleveland Clinic and the Case Western Reserve University Genomics Core for their services; the Pathology Research Core in the Pathology and Laboratory Medicine Institute of Cleveland Clinic and the Clinical Core of Cleveland Clinic’s Northern Ohio Alcohol Center (funded by NIH grant P50AA024333) for their human tissue services; the Howard Hughes Medical Institute Janelia GENIE project for developing and sharing the CaMPARI sensor; and current and prior members of the Davalos and Nagy laboratories for discussions and experimental assistance. This work was supported by a Bodossaki Foundation Postdoctoral Fellowship (E.P.); a Cleveland Clinic Research Co-Laboratories grant (D.D. and L.E.N.); the NIH National Institute on Drug Abuse award P30 DA054557 (S.V.); the NIH BRAIN Initiative award U01NS123658 (H.D.); the National Institute for Neurological Disorders and Stroke award R01NS112526 (D.D.); the NIH National Institute on Alcohol Abuse and Alcoholism awards U01 AA029969 (S.D.B.) and P50AA024333 (L.E.N. and D.D.). The content is solely the responsibility of the authors and does not necessarily represent the official views of their employers or the National Institutes of Health. Illustrations were made using BioRender.

## Methods

### Animal models and treatments

Mice were bred in the Biological Resources Unit at Cleveland Clinic Research under standard laboratory conditions (20–22°C, 30–70% humidity, 12-hour light/dark cycle, ad libitum food and water). All experiments were performed under Cleveland Clinic–approved Institutional Animal Care and Use Committee protocols. Mouse strains acquired from the Jackson Laboratories: C57BL/6J (000664), *Cx3cr1*^GFP/GFP^ (005582), *MyD88*^F/F^ (008888), *Rosa26*^tdTomato^ (007914), *Cx3cr1*^creERT2^ (020940), *Tmem119*^creERT2^ (031820), *MyD88*^−/−^ (009088). Dr. Staci Bilbo provided the *Cx3cr1*^creTg/0^ (MGI:5311737) mouse line. These lines were then bred to generate all conditional knockout and reporter lines used in the study. Genotyping was performed by PCR analysis of tail DNA. *Cx3cr1*^creERT2^ mice have high recombination efficiency in microglia but also target brain perivascular/meningeal macrophages and some peripheral macrophages^94^, while *Tmem119*^creERT2^ mice have lower efficiency but higher microglial specificity^95^. Ethanol experiments began ≥4–5 weeks after tamoxifen induction to ensure specific microglial targeting while allowing replenishment of recombined blood monocytes and tissue-resident macrophages^94^. Both sexes were used in Supp. Fig. 1a, b; no sex differences in ethanol-induced microglial reactivity were observed, so male mice were used thereafter. Experimental mice were 14–20 weeks old.

#### Ethanol administration: Repeated binge drinking models

Moderate-to-heavy episodic (“binge”) drinking is the most common form of alcohol abuse in the United States. To simulate a pattern of repetitive binge or high intensity alcohol intake, adult mice (14–20 weeks old) received daily oral gavage of EtOH (binge: 2 g/kg BW: 25.3% v/v, or high intensity: 5 g/kg BW: 63.4% v/v in 0.9% saline) for up to 10 consecutive days. Control mice received equal volumes of 0.9% sterile saline. After ethanol administration, cages were positioned 50% over heating pads until recovery, to minimize hypothermic effects of ethanol. Mice were euthanized 3 or 5–6 hours after final gavage (recovery from ethanol-induced ataxia (binge dose) or sedation (high-intensity dose), respectively) and tissues were collected acutely for downstream analyses.

#### Tamoxifen induction

Tamoxifen-inducible mice received five consecutive daily intraperitoneal injections of 2 mg tamoxifen in corn oil at 6–10 weeks of age, ≥4–5 weeks before ethanol experiments.

### Blood ethanol measurements

Mice orally received ethanol (in saline) and were returned to home cages for 90 min (half cage over heating pad). Tail blood was collected in hematocrit capillary tubes (Fisher), and plasma ethanol concentration was measured spectrophotometrically at 340 nm using the Ethanol L3K assay (Sekisui Diagnostics) according to the manufacturer’s instructions and as previously^96^.

### *In vivo* imaging using two-photon microscopy and image analyses

#### Cranial window implantation

Mice were anesthetized with isoflurane (2.5–3% induction, 1.5– 2% maintenance) and maintained at 37°C. After stereotaxic head-fixation, aseptic surgery, and bupivacaine injection (2 mg/kg, SC), a ∼3 mm craniotomy was performed over the right somatosensory cortex. The craniotomy was sealed with glass coverslips (two 3 mm and one 5 mm in diameter, Warner Instruments), two 3 mm and one 5 mm in diameter, glued together using UV-cured optical adhesive (Norland Products). The cranial window was secured with cyanoacrylate glue, and a metallic head bar was attached using dental cement. Post-operative analgesia included: buprenorphine SR (3.25 mg/kg, SC) and ketoprofen (1.5 mg/kg, SC daily for 48 h). For neuronal activity recordings, adeno-associated virus (AAV) encoding the CaMPARI1 calcium indicator under the human synapsin promoter (php.N-hSyn-CaMPARI1, Molecular Tools Platforms, CERVO Brain Research Centre; 40 μL, ∼1×10¹³ GC/ml) was injected retro-orbitally ∼1 week post-surgery. Mice were allowed a minimum recovery time of 21 days before imaging experiments began.

#### Two-photon microscopy for microglial imaging

Two custom-built two-photon laser-scanning microscopes (Ultima IV and Investigator, Bruker) equipped with Chameleon Vision II and Discovery lasers (Coherent) were used for *in vivo* imaging of microglia. Excitation: 880–920 nm for EGFP and vascular labels; 1040 nm for tdTomato. Mice were habituated to head-fixation under the two-photon microscope using a treadmill (Luigs & Neumann) for ≥3 days. Imaging was performed in Layer I (80–100 μm depth) using Nikon 16× 0.8 NA or 25× 1.1 NA lenses with 512×512 pixel resolution and 2.48–3.7× zoom. Z-stacks (80 μm depth, 1 μm z-step) acquired over ∼45 seconds for 2–3 cortical regions per mouse. Blood vessels were labeled with retro-orbital injection of Rhodamine B or Fluorescein Dextran (ThermoFisher Scientific) for field relocation. Imaging sessions: Daily imaging included baseline (pre-gavage) and 1, 2, 3–4, 5–6 hours post-gavage, and was repeated on saline day and ethanol (5 g/kg BW) days 1, 3, 5, 10 for each animal. The imaging chamber was maintained at 37°C during EtOH exposure imaging sessions.

#### Two-photon microscopy for neuronal activity recordings

Following a ∼4-week surgical recovery and viral expression period, mice received saline or ethanol (5 g/kg BW), then 10-min photoconversion 1-2 hours post-gavage using a 405 nm LED (UHP-X-405, Prizmatix; 13.5 W focused to ∼0.1 mW/mm²). Then, the mice were placed under a custom-built two-photon microscope using galvo/resonant scanners (MiMMS 2.0, designed by HHMI Janelia Research Campus). Simultaneous recording of green and red CaMPARI1 signals was performed using 1000 nm excitation light (Insight X3, Spectra-Physics), with a 16× 0.8 NA objective lens (Nikon). Images were captured at 15 frames per second with 1024×1024 pixels per image covering an area of 1.2 × 1.2 mm. Each cortical column (z-stack) consisted of 6 distinct z-planes, 45μm apart, spanning 45–270 μm from the pial surface. 3-7 columns, spanning the somatosensory cortex in one hemisphere, were recorded from each mouse and were used for the analysis. The same animals and brain locations were reimaged with 9-10-day intervals (saline day, and ethanol days 1, 10).

#### Microglial morphology analyses

Z-stacks of images spanning 80 μm in depth were acquired, corrected for focal plane displacement using ImageJ (NIH) with TurboReg and StackReg plug-ins, and trimmed to relocate the same microglial cells within a z-depth of 50 μm across all timepoints. Analyses were performed in Imaris 10.2.0 (Oxford Instruments) using the Filaments module, after background subtraction. Perivascular macrophages were excluded. Surveillance territory was normalized to the average value from all imaged locations recorded during the saline treatment day for each animal. Surveillance territory and soma volume changes were calculated as percent of corresponding values averaged from images collected during the saline day recordings.

#### Neuronal activity analysis

Custom MATLAB (Mathworks) scripts were used for analyzing CaMPARI signals from individual neuronal cell bodies. Cell bodies were segmented using CellPose2^97, 98^ first automatically, and then additional cells were segmented manually to ensure accurate sampling across conditions. Bleed-through light contamination of the green fluorescence into the red channel was corrected using data from pre-photoconversion recordings^65^. Images with less than 10 detected somata were excluded from the analysis. Small diameter red puncta (noise) could be identified in the red channel and were segmented and excluded from the analysis prior to red to green (RG) ratio calculations using the Matlab Image Segmenter App. Missing animal datapoints in Fig. 5e are due to animal death (one wild-type and one conditional knockout) before ethanol day 10.

### Histology, immunofluorescence and analyses

#### Tissue collection

Mice were anesthetized with ketamine/xylazine (100–200/15–30 mg/kg BW), blood was collected from the vena cava into EDTA-coated tubes (BD Microtainer), then transcardially perfused with ice-cold phosphate-buffered saline (PBS). Brains were fixed in 4% paraformaldehyde (PFA) in PBS (48 h, 4°C), cryoprotected in 30% sucrose, embedded in OCT (Tissue-Tek), and sectioned sagittally (50 μm for microglial analyses, 30 μm for other histology) on a cryostat (Leica Microsystems). Serial free-floating sections were stored in cryoprotectant (30% ethylene glycol, 20% glycerol in PBS) at −20°C. Liver tissue and isolated plasma were snap frozen in liquid nitrogen.

#### Mouse brain immunofluorescence

Sections were washed in PBS (3×5 min), permeabilized with 1% Triton-X in PBS (45 min for Homer1/SYP only), blocked with 5% normal donkey serum (NDS, SouthernBiotech), 3% bovine serum albumin (BSA, Sigma-Aldrich) 0.3% Triton-X in PBS for 1 h at room temperature (RT), incubated with primary antibodies in 1% BSA, 0.3% Triton-X in PBS, overnight at 4°C, washed, incubated with secondary antibodies (1–1.5 h), washed, and mounted with DAPI medium (Aqueous Fluoroshield, Abcam).

#### Microscopy and quantification

Imaging was performed on anatomically equivalent areas for all experimental comparisons, using LSM800 (Zeiss) or Stellaris 5 (Leica) confocal microscopes with the ZEN3.7 or LAS X software, respectively. 40× 1.3 NA (microglia, CD68, PSD95) or 63×1.4 NA oil-immersion lenses (Homer1/synaptophysin, microglia/CD68, HA). Imaris 10.2.0 was used for 3D analyses: Filaments module for 3D tracing of individual microglia and their soma volume and process length, branching, volume, and surveillance territory calculations; Surfaces module for 3D rendering and quantification of microglial lysosomes and synaptic marker quantification. Homer1-masked synaptophysin was reported as synapses; microglia-masked CD68 as microglial lysosomes; microglial CD68-masked PSD95 as engulfed material. The Gaussian filter and background subtraction in Imaris were used to reduce noise. For NeuN, z-stack images were acquired using a 10× 0.45 NA or 20× 0.75 NA lens under a BZ-X710 epifluorescence microscope (Keyence) with the BZ-X700 Viewer software. Quantification of NeuN+ cells was performed using either the Hybrid Cell Counter of the BZ-X Analyzer software or the Spots module in Imaris. 1 image per section from 2–3 sections per mouse brain region were used for analyses.

### Cell isolation and molecular analyses

#### FACS isolation of microglia

Following ketamine/xylazine anesthesia, mice were perfused with ice-cold PBS and whole brains were dissected minced and digested for generation of single-cell suspensions (dispase II 10 mg/ml, collagenase VIII 1.4 U/ml, DNAse I 50 U/ml, 5 mM CaCl₂, 30 min, 37°C, 120 rpm). For RNA-sequencing experiments, transcription and translation inhibitors were added to perfusion, tissue collection, and enzyme solutions as previously described^99^. After serial pipette triturations, filtering through 70 μm cell strainer, and washing with 20% fetal bovine serum (FBS), 25mM HEPES in RPMI1640, cells were separated from myelin and other tissue debris on 30% Percoll gradient (GE Healthcare) at 800g for 30min (half acceleration, no brake), all at 4°C. Cells were resuspended in RPMI1640 containing 25 mM HEPES buffer and 1% FBS, and labeled with antibodies for 30min and DAPI (to identify dead cells). DAPI⁻ CD45^int^ CD11b⁺ microglia were collected on a FACSAria Fusion cell sorter (BD Biosciences) using a 100 μm nozzle with purity mode. Unstained or DAPI-labeled lymph node cells and single-stained OneComp eBeads Compensation Beads (ThermoFisher Scientific) were used for compensation settings.

#### qPCR for MyD88 recombination

After cell sorting, microglial genomic DNA was extracted from pelleted cells using the DNeasy Blood & Tissue Kit (Qiagen), and ∼50 ng was used for qPCR with published primers and cycling conditions for the detection of the functional unrecombined –but not the null– *MyD88* allele^62^. PowerUp SYBR Green Master Mix 2X (ThermoFisher Scientific) and 0.5µM of each primer was used. Cycling was performed in 384-well plates (Greiner Bio-One) on a QuantStudio 6 Flex instrument (Applied Biosystems) using QuantStudio™ Real-Time PCR Software 1.7.2. For data normalization, the Ct value of *k17* was determined as the reference for each sample^62^, and the relative quantification method (2^-ΔΔCt^) was used.

#### RNA sequencing

Murine microglia were isolated from the brains of *Cx3cr1^creERT2^:Rosa26^tdTomato^* mice 18 hours after the final of four daily gavages with 5 g/kg BW EtOH. Due to technical limitations, five of the six microglia samples from saline-treated control mice were isolated and sequenced in a separate batch (after 10 days of saline treatment). Microglial cells were directly sorted into RLT Plus lysis buffer, and total RNA was extracted using the RNeasy Plus Micro Kit (Qiagen). RNA sample concentrations and RIN scores were determined by the Bioanalyzer RNA 6000 pico assay (Agilent). A total-mRNA library was prepared using the SMART-Seq mRNA LP (with UMIs) kit (Takara Bio USA). This involved oligo(dT) priming for polyA transcript selection and a 100 bp fragment size selection. RNA sequencing was performed on the NovaSeq X platform (Illumina), yielding 72M-106M paired-end reads per sample. Quality control was performed with FastQC v0.12.1 and MultiQC v1.19^100, 101^. Sequencing primers and adapters were trimmed using Bbduk v39.01^102^, and trimmed sequences were re-assessed with FastQC v0.12.1^100^. Trimmed reads were aligned to the GRCm39 mouse genome using Hisat v2.2.1^103^, with 85–90% of reads successfully mapped. Rsubread v2.14.2^104^ was used for quantifying gene expression values. Differential expression tests between experimental groups (EtOH vs. Saline) were performed using DESeq2 v1.40.2^105^, with batch included as a covariate in the design formula (design = ∼ batch + condition) to correct for batch effects. Genes with an adjusted P < 0.05 and an absolute |Log2Fold Change| > 0.6 were considered statistically significant. Pathway analysis was conducted using GSEA with the pre-ranked option using mSigDB gene lists, including Hallmark, KEGG, Reactome, and Gene Ontology (GO).

#### Isolation of liver RNA and qRT-PCR

Total RNA was isolated from liver tissue using Rneasy Plus Universal Mini Kit (Qiagen) and reverse transcribed with SuperScript™ IV VILO™ Master Mix (Invitrogen), followed by amplification using qRT-PCR. Power SYBR^TM^ Green PCR Master Mix (Applied Biosystems) and primers (listed below) at final concentrations of 1 μM were used. Cycling was performed in a QuantStudio5 instrument (Applied Biosystems) for 40 cycles of 15 s at 95°C, 30 s at 60°C, 30 s at 72°C followed by 1 min at 95°C, 30 s at 55°C and 30 s at 95°C. Data were analyzed using the 2^-ΔΔCt^ method, with 18S as the reference gene.

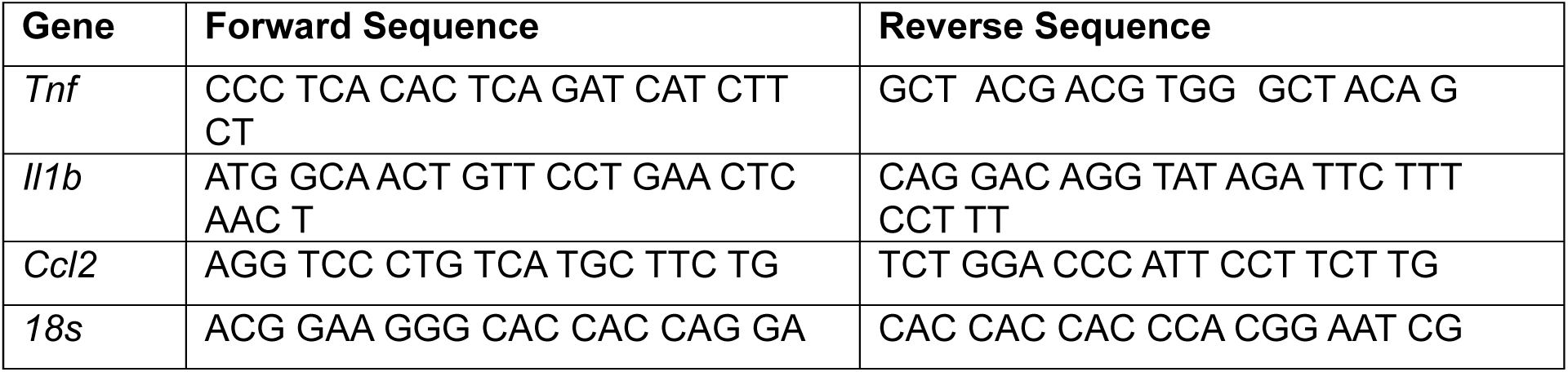

### Reagents

#### Antibodies

The following antibodies were used in this study: Iba1 (CP290B, BioCare Medical, 1:200; 019-19741, Wako, 1:1000), NeuN (MAB377, Millipore, 1:500), Homer1 (160002, Synaptic Systems, 1:200), Synaptophysin (sc-17750, Santa Cruz Biotechnology, 1:200), PSD-95 (MAB1596, Millipore, 1:100), CD68 (MCA1957, Biorad, 1:250). Secondary antibodies: Cy5-conjugated anti-rabbit (711-175-152, Jackson ImmunoResearch, 1:200), Cy5-conjugated anti-mouse (715-175-151, Jackson ImmunoResearch, 1:100), Cy5-conjugated anti-rat (712-175-150, Jackson ImmunoResearch, 1:400), Cy3-conjugated anti-mouse (715-165-150, Jackson ImmunoResearch, 1:1000), Cy3-conjugated anti-rabbit (711-165-152, Jackson ImmunoResearch, 1:1000), FITC-conjugated anti-rabbit (711-095-152, Jackson ImmunoResearch, 1:1000). Flow cytometry: anti-CD16/32 (553142, BD Biosciences, 1:100), PECy7-conjugated anti-CD45 (552848, BD Biosciences, 1:100), FITC-conjugated CD11b (101205, Biolegend, 1:100).

#### Biochemical assays

Commercially available kits were used for the quantification of plasma ALT/AST activity (Sekisui Diagnostics), and total liver triglycerides (Pointe Scientific) per manufacturers’ instructions. For alcohol dehydrogenase (ADH) activity, whole liver was homogenized in Lysing Matrix D tube (MP Biomedicals) in the ADH buffer provided in the ADH assay kit and activity quantified using a colorimetric enzymatic assay (Sigma Aldrich) per the manufacturer’s instructions.

### Behavioral tests

#### Rotarod

This test was used to assess sensorimotor coordination and motor learning. Mice were placed on a non-accelerating rotating rod (fixed speed at 16 rpm, 3 cm diameter rod, 44.5 cm height, Rotamex-5; Columbus Instruments). Each mouse was trained until able to remain on the rotarod for 180 s, one day prior to experimental testing. The latency to fall was detected by an infrared beam sensor and recorded as the duration that mice remained on the rod, with a cutoff latency of 180 s. On the experimental testing day, mice underwent reminder/retraining until they could stay on the rod for 60 s, then gavaged with 0.9% saline or ethanol (2 g/kg BW) and placed back on the rotarod 15 minutes later. Latency to fall was measured every 15 minutes until each mouse was able to remain on the rotarod for 60 s. Any animal able to remain on the rotarod for 60 s during the first trial following ethanol administration (15 min), was tested again for a second trial at 30 minutes.

#### Locomotor activity

This test was used to assess locomotor activity over time in a home-cage environment (width 21 cm × length 43 cm × height 29 cm with clean minimal bedding and a cage lid) and was performed under red light to maximize the amount of mouse activity. Clean home cages were positioned between stacked infrared photobeam frames (SD Instruments PAS-HC). X-Y beams (3 cm height) were used for detection of ambulatory activity, Z beams (8 cm height) for rearing. Up to twelve individually housed mice were tested at once, beginning with 1 h habituation to the cage, followed by oral gavage (0.9% saline or ethanol 2 g/kg BW) and 2-hour recording of activity, which was reported as number of beam breaks in 5 min time bins.

#### Loss of righting reflex (LORR)

This test was used to assess the sedative effects of alcohol as previously^44, 60^. Mice were injected intraperitoneally with ethanol (3.5 g/kg BW, 19.7% v/v in saline). When mice became ataxic and completely stopped moving in their home cage, they were placed supine in plastic V-shaped troughs until they could right themselves 3 times in 30 seconds. Latency to LORR: time elapsed from ethanol administration to being unable to right themselves within 30 s from supine placement. Duration of LORR: Time elapsed from LORR to three successful rightings within 30 s. Body temperature was maintained with heat lamp.

### Statistical analyses

All measurements were taken from distinct samples. Statistical analyses were performed using GraphPad Prism v.8.1.1 (GraphPad Software Inc.). Data presented as mean ± SEM assuming normal distribution. Sample sizes, statistical tests, and p-values are reported in figures and figure legends.

