## Supplementary Figures for "Preventing Microglial Reactivity Protects from Acute and Progressive Neuronal Dysfunction, Motor Impairments and Sedation following Alcohol Abuse"

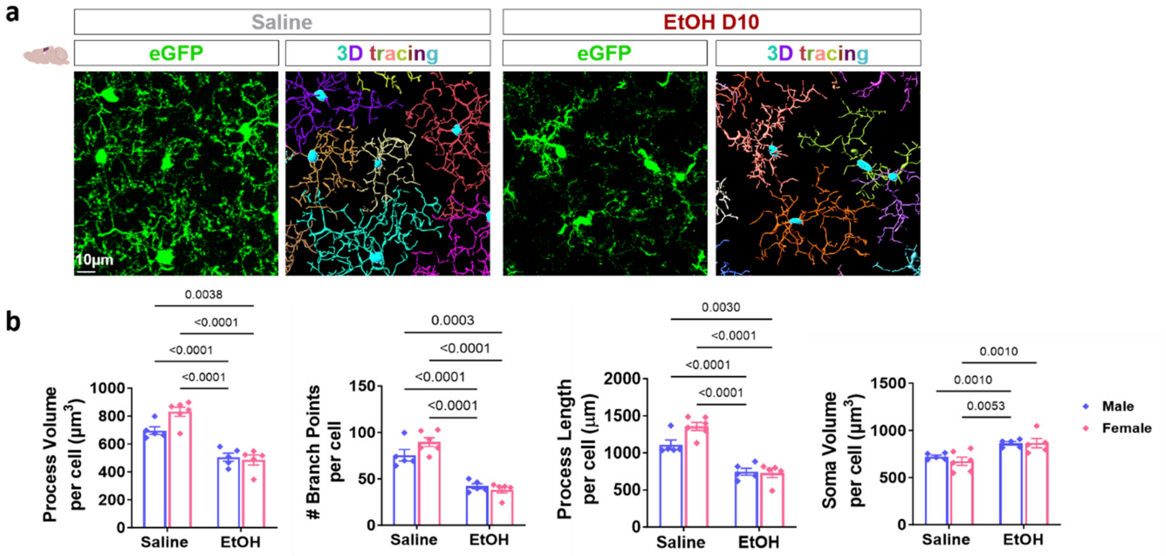


c

| Alcohol Intoxication Scoring System | |
|---|---|
| 0 | Normal behavior |
| 1 | Lethargic, dizzy, wobbling when walking |
| 2 | Laying down, head still moving and alert, will walk/move if disturbed |
| 3 | Laying down, appears unconscious, responds to cage tapping |
| 4 | Fully unconscious, does not respond to cage tapping |
| 5 | Dead |

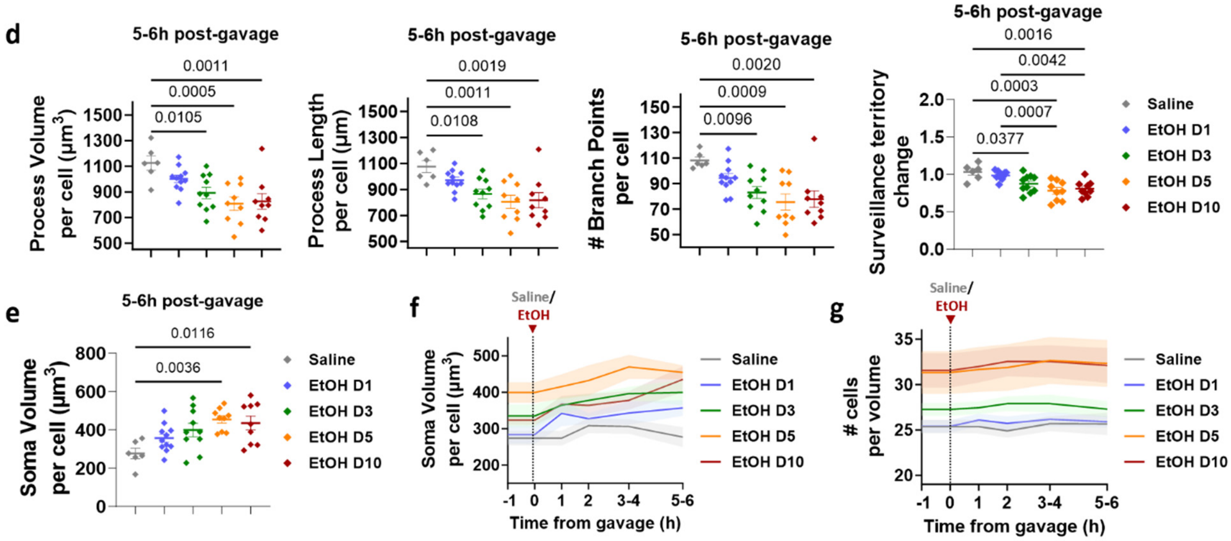


**Supp. Fig. 1. Repeated binge ethanol consumption induces progressive morphological microglial reactivity, that is similar in males and females. a.** Representative confocal microscopy images of microglia and 3D tracing of their processes and cell body reconstruction in somatosensory cortex of *Cx3cr1*$^{GFP/+}$mice. Images were taken from cortical tissues collected from mice on day 10 of their treatment with saline or ethanol, as described in Fig. **1a**(**i**). Scale bar, 10 μm. **b.** Quantification of microglial process volume, branch points, process length, and soma volume in somatosensory cortex of male and female *Cx3cr1*$^{GFP/+}$ mice after 10 consecutive days of treatment as described in Fig. **1a**(**i**). n = 5 mice per group; Two-way ANOVA with Tukey's multiple comparison test. **c.** Alcohol intoxication scoring system developed by observation of symptoms upon ethanol oral administration as described in Fig. **1a**(**i**), similar to a prior study[58]. **d-e.** Quantification of microglial process volume, length, branch points, surveillance territory change **(d)** and soma volume **(e)** at 5–6 h post-gavage on saline day and binge ethanol exposure days 1, 3, 5, and 10. One-way ANOVA with Tukey's multiple comparison test. **f-g.** Time course changes of microglial soma volume **(f)** and cell number **(g)** per analyzed volume over hours following gavage on saline day and on binge ethanol exposure days 1, 3, 5, and 10. Data presented as mean ± SEM. **d-g.** n = 11 imaged locations from 6 *Cx3cr1*$^{GFP/+}$ mice. Only statistically significant comparisons are shown.

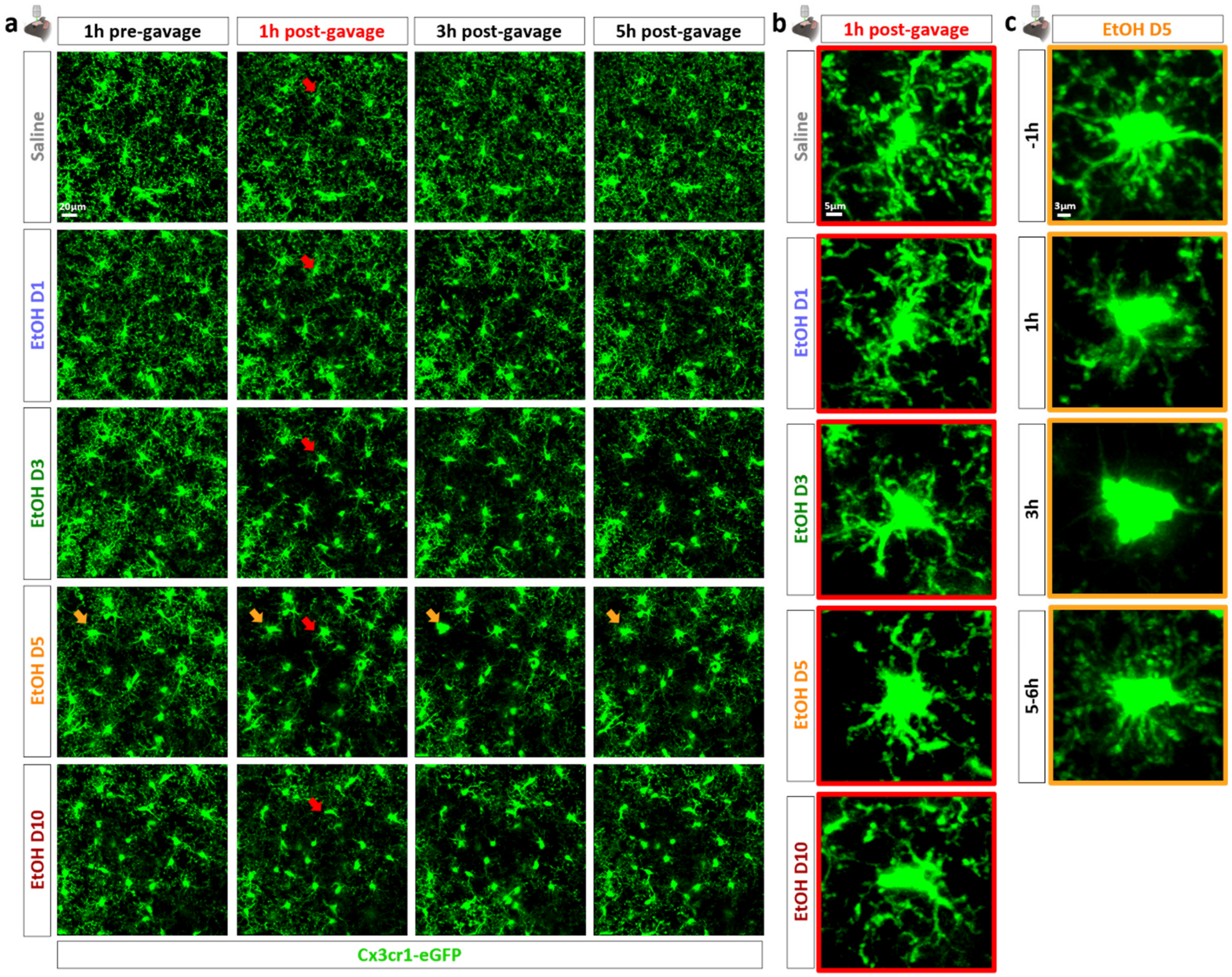


**Supp. Fig. 2. Repeated ethanol abuse induces acute and progressive morphological microglial responses *in vivo.* a.** Representative maximum intensity projections of 50 µm z-stacks used for 3D morphological analysis of cortical microglia imaged with two-photon microscopy *in vivo* in real time. Images show the same cortical location at -1 h, +1 h, +3 h, and +5–6 h following gavage on saline day and on binge ethanol exposure days 1, 3, 5, and 10. Scale bar, 20 µm. **b.** Morphological changes of the processes and soma of a single microglial cell (indicated with red arrows in panel **a**) at 1 h post-gavage on saline day and on binge ethanol exposure days 1, 3, 5, and 10. Scale bar, 5 µm. **c.** Example of a microglial cell with robust loss of ramification and enlargement of cell body over time on ethanol day 5 of the 10-day binge drinking course (indicated with orange arrows in panel **a**). Scale bar, 3 µm.

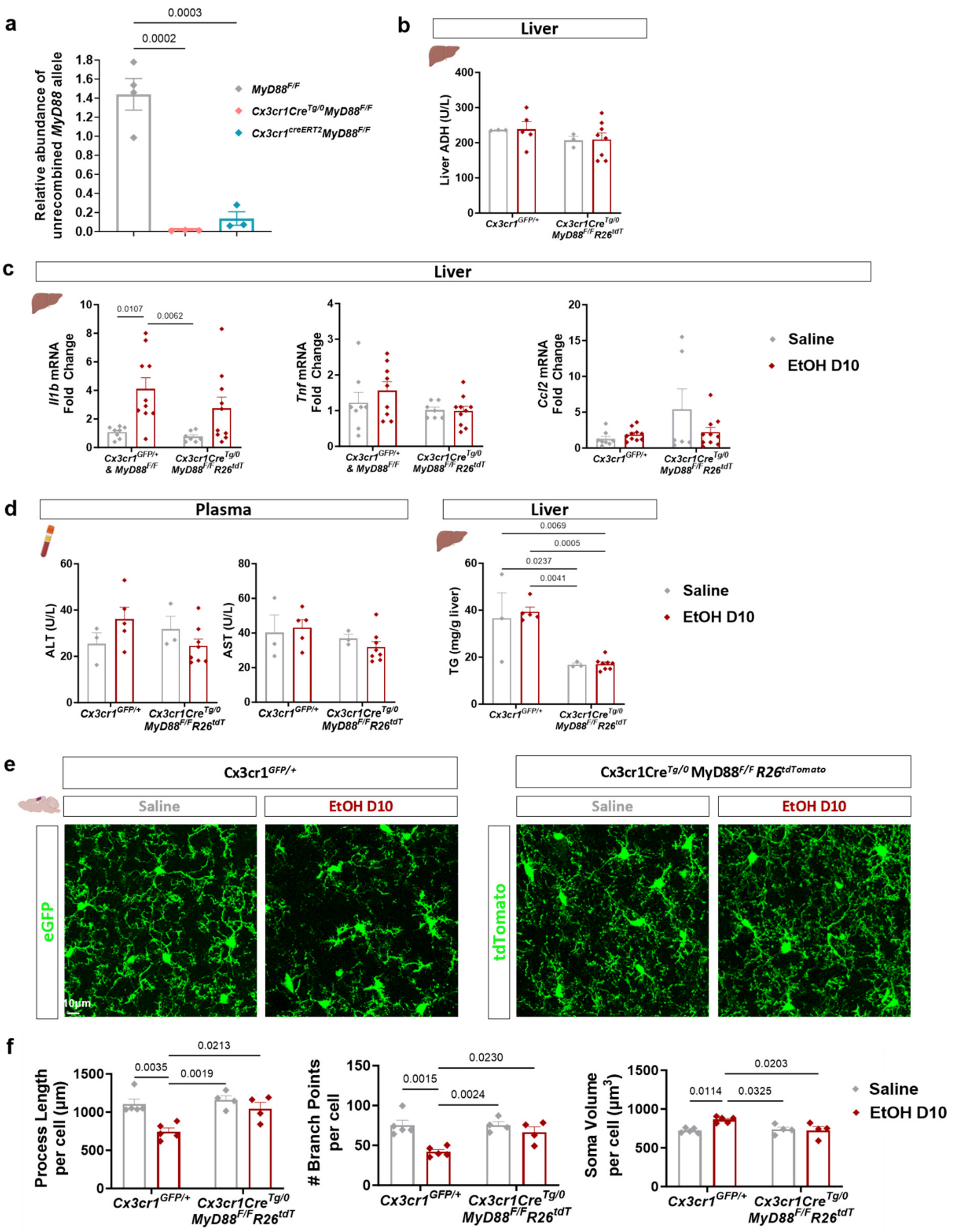


**Supp. Fig. 3. Macrophage-specific deletion of *MyD88* protects from ethanol-induced microglial activation, without affecting ethanol metabolism and liver inflammation. a.** qPCR for the unrecombined *MyD88* allele in genomic DNA isolated from FACS-sorted microglia from *MyD88*$^{F/F}$, *Cx3cr1cre*$^{Tg/0}$;*MyD88*$^{F/F}$ and *Cx3cr1cre*$^{creERT2}$;*MyD88*$^{F/F}$ mice. n = 3–4 mice per group;

One-way ANOVA with Tukey's multiple comparison test. **b.** Quantification of liver alcohol dehydrogenase (ADH) in saline- and ethanol-treated *Cx3cr1*$^{GFP/+}$ and *Cx3cr1cre*$^{Tg/0}$*;MyD88*$^{F/F}$*;Rosa26*$^{tdTomato}$ mice on day 10 of binge ethanol exposure (5 g/kg BW). n = 3-8 mice per group. **c.** qPCR for the detection of *Il1b*, *Tnf* and *Ccl2* mRNA expression in livers from mice treated as described in (**b**). n = 7-10 mice per group. **d.** Quantification of plasma alanine transaminase (ALT) and aspartate transferase (AST) and liver triglycerides (TG), in mice treated as described in (**b**). n = 3–8 mice per group. **e.** Representative confocal microscopy images of eGFP- and tdTomato-labeled microglia in somatosensory cortex of mice treated as described in (**b**). Scale bar, 10 µm. **f.** Quantification of microglial process length, branch points, and cell body volume in somatosensory cortex of mice treated as described in (**b**). n = 4-5 mice per group. Data presented as mean ± SEM. **b-d, f:** Two-way ANOVA with Tukey's multiple comparison test. Only statistically significant comparisons are shown.

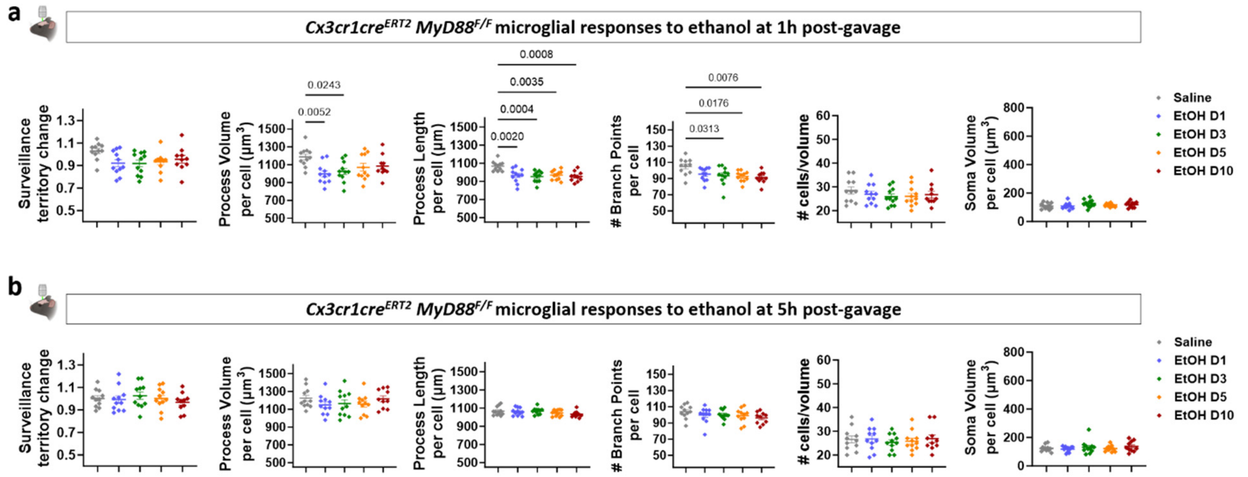


**Supp. Fig. 4. Microglial-specific *MyD88* deletion prevents ethanol-induced morphological changes of microglia over time *in vivo.* a, b.** Quantification of microglial surveillance territory change, process volume, process length, branch points, cell density and soma volume, in *Cx3cr1*$^{creERT2}$;*MyD88*$^{F/F}$;*Rosa26*$^{tdTomato}$ mice at 1 h **(a)** and 5–6 h **(b)** post-gavage on saline day and binge ethanol exposure days 1, 3, 5, and 10. Data presented as mean ± SEM. n = 11 imaged locations from 6 mice. One-way ANOVA with Tukey's multiple comparison test. Only statistically significant comparisons are shown.

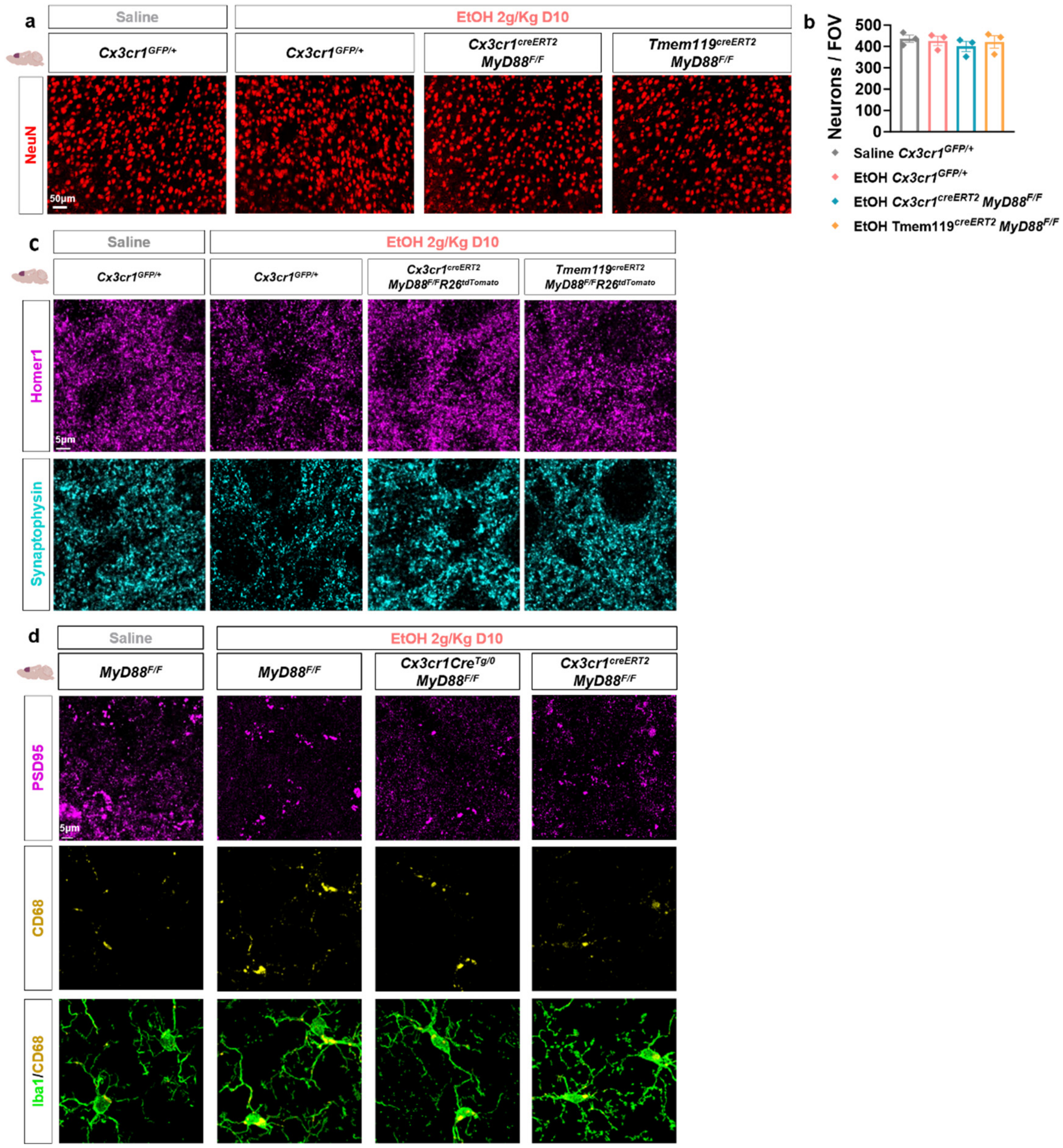


**Supp. Fig. 5. Microglia-specific *MyD88* deletion protects from synapse loss and increase of CD68⁺ lysosomes in microglia. a.** Representative epifluorescence microscopy images of NeuN-labeled neurons in prefrontal cortex of saline- and ethanol-treated mice on day 10 of binge ethanol exposure (2 g/kg BW). Scale bar, 50 µm. **b.** Quantification of NeuN⁺ neurons per field of view (FOV) in prefrontal cortex of mice treated as described in (**a**). Data presented as mean ± SEM. n = 3 mice per group. One-way ANOVA with Tukey's multiple comparison test. **c.** Individual channels (related to Fig. **4a, b**) for post-synaptic Homer1 and pre-synaptic synaptophysin labeled by immunofluorescence in prefrontal cortex of mice treated as described in (**a**). Scale bar, 5 µm. **d.** Representative confocal microscopy images (related to Fig. **4f, g**) of immunolabeled Iba1⁺ microglia, CD68⁺ lysosomes and PSD-95⁺ post-synaptic densities in prefrontal cortex of mice treated as described in (**a**). Scale bar, 5 µm.

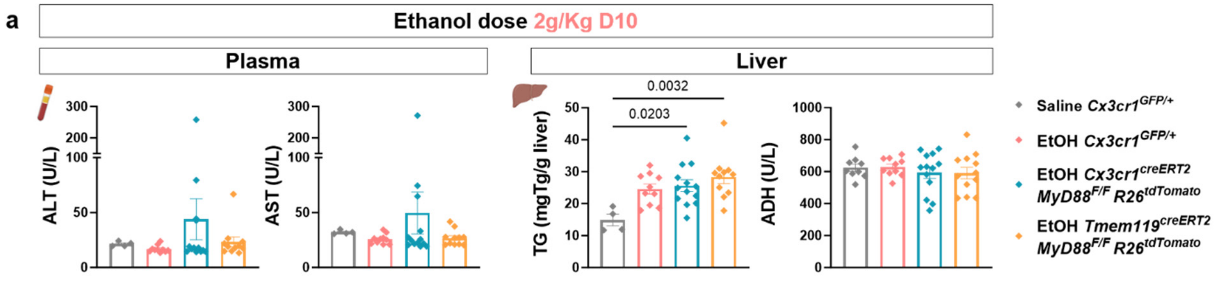


**Supp. Fig. 6. Microglia-specific *MyD88* deletion does not alter ethanol metabolism and liver damage markers. a.** Quantification of plasma alanine transaminase (ALT), aspartate transferase (AST), liver triglycerides (TG) and alcohol dehydrogenase (ADH) in saline- and ethanol-treated mice on day 10 of binge ethanol exposure (2 g / kg BW). Data presented as mean ± SEM. n = 4–13 mice per group. One-way ANOVA with Tukey's multiple comparison test.
